# A dynamical analysis of the alignment mechanism between two interacting cells

**DOI:** 10.1101/2024.07.23.604626

**Authors:** Vivienne Leech, Mohit P. Dalwadi, Angelika Manhart

## Abstract

In this work we analytically investigate the alignment mechanism of self-propelled ellipse-shaped cells in two spatial dimensions interacting via overlap avoidance. By considering a two-cell system and imposing certain symmetries, we obtain an analytically tractable dynamical system, which we mathematically analyse in detail. We find that for elongated cells there is a half-stable steady state corresponding to perfect alignment between the cells. Whether cells move towards this state (i.e. become perfectly aligned) or not is determined by where in state space the initial condition lies. We find that a separatrix splits the state space into two regions, which characterise these two different outcomes. We find that some self-propulsion is necessary to achieve perfect alignment, however too much self-propulsion hinders alignment. Analysing the effect of small amounts of self-propulsion offers an insight into the timescales at play when a trajectory is moving towards the point of perfect alignment. We find that the two cells initially move apart to avoid overlap over a fast timescale, and then the presence of self-propulsion causes them to move towards a configuration of perfect alignment over a much slower timescale. Overall, our analysis highlights how the interaction between self-propulsion and overlap avoidance is sufficient to generate alignment.

## 1 Introduction

### Alignment in biology

Alignment of particles in a system is a phenomenon that can be observed in many contexts. In biology, this ranges from the large scale e.g. schools of fish (Herbert-Read et al., 2011), down to the microscopic scale e.g. cells and bacteria (Balagam and Igoshin, 2015; Dartsch and Betz, 1989; Kenny et al., 2023). This alignment of particles, especially when it occurs collectively, can play key roles in their migration e.g. hydrodynamic benefits as a result of collective alignment can aid migration in schools of fish (Lopez et al., 2012) and alignment of fibroblasts can affect key mechanical properties of the tissue in which they are found (Erdogan et al., 2017).

### Motivation and context of this work

Many collective alignment models (Peruani et al., 2006; Baskaran and Marchetti, 2008; Kraikivski et al., 2006) include self-propulsion and some kind of overlap avoidance or volume exclusion as model ingredients, suggesting that these are key components needed for collective alignment between particles to occur. This can be understood intuitively, since overlap avoidance provides a way for cells to change their orientation in reaction to other cells while self-propulsion ensures that cells continue to interact with each other, allowing for alignment to propagate through the population. Motivated by experiments on the alignment of fibroblasts in Kenny et al. (2023), an agent-based model was recently developed in Leech et al. (2024) to investigate the mechanism behind the collective alignment of self-propelled interacting particles. The model is set in two spatial dimensions and describes a collective of ellipse-shaped cells with fixed area. These cells move in the direction of their orientation and change their position, orientation and shape in order to avoid overlap. In Leech et al. (2024) and also in this work, we use the term *overlap avoidance* rather than volume exclusion to emphasis that overlap is allowed in principle, but punished by a tunable potential. Cell overlap then corresponds to cells being positioned partly on top of each other, an observed phenomena for cells crawling on surfaces (see a 3D interpretation in Fig. 1B). Through computational analysis of the model, it is found that these model components lead to collective alignment, with the amount and spatial scale of the alignment depending on model parameters.

**Figure 1:**
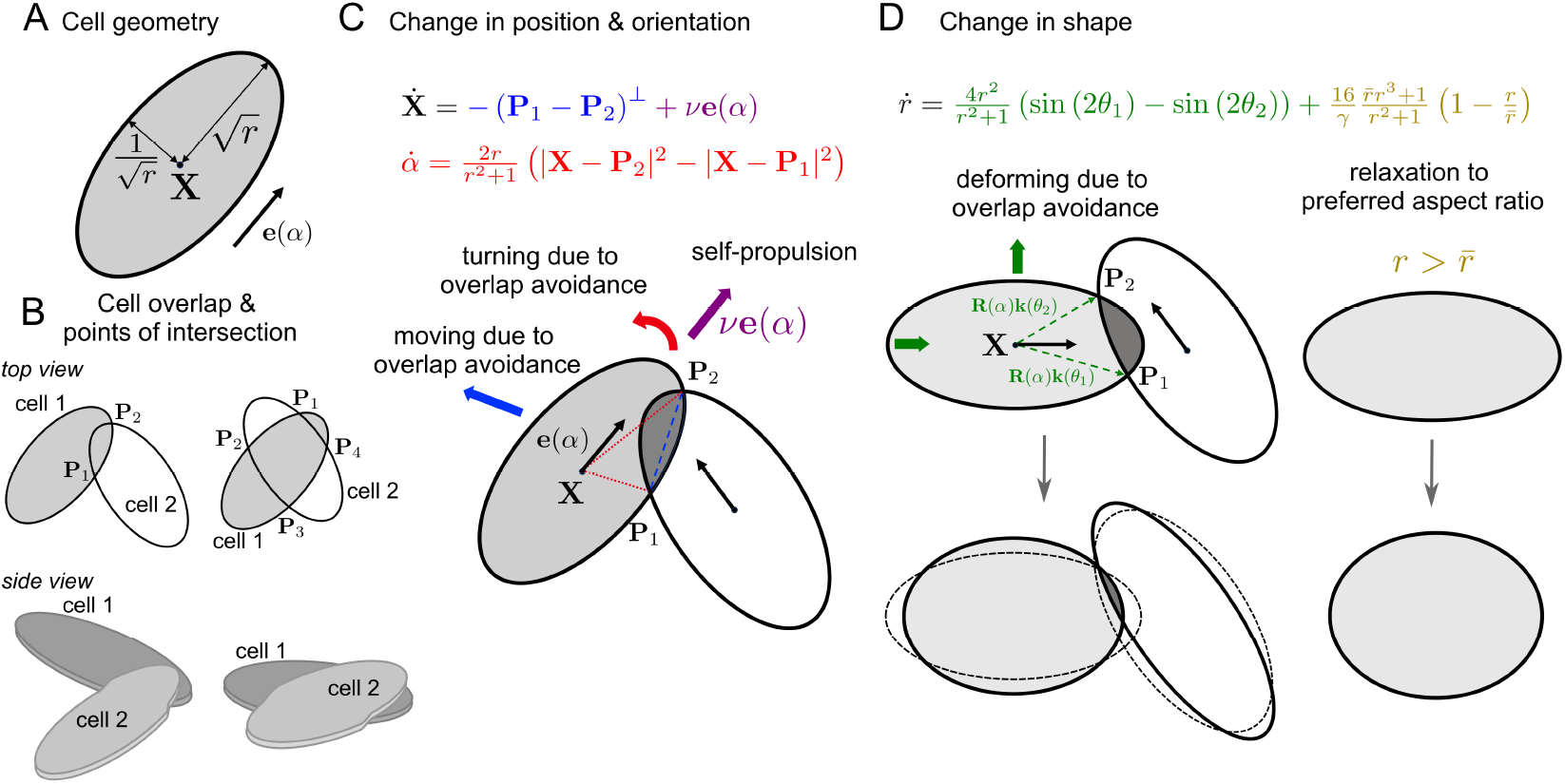
A: (Non-dimensional) cell geometry. B: Naming of intersection points. C, D: Equations and schematic for interaction of two cells for changes in position & orientation (C) and aspect ratio (D).

### Limitations of this work

There are several limitations of this work. Firstly, it does not capture collective effects, however by comparing to Leech et al. (2024) we can learn which phenomena are likely consequences of the underlying alignment mechanisms and which are driven by collectivity. Secondly, the symmertry condition imposed in this work limits *alignment* to “velocity alignment” (where cells move in the same direction) and cannot capture “nematic alignment” (where cells might also move in opposite directions). Also, not all model ingredients of Leech et al. (2024) are included in this analysis, most notably we omitted cell-cell adhesions and any cytoskeletal forces transmitted through them (see also the discussion at the end of this work). Finally, the model of Leech et al. (2024) itself already omits various biological mechanisms that have been shown to affect alignment, at least in some situations. Examples are feedback with the substrate (Wang et al. (2018)), the effect of a surrounding fluid (Ng and Swartz (2003)) or more complicated cell signalling.

### The challenge of analysing agent-based model

The most common approach for the analysis of agent-based models tends to be computational (as was the case in Leech et al. (2024)), since there are fewer analytic tools for analysing agent-based models. For an overview of different approaches to modelling and analysing pattern formation, see e.g. Deutsch and Dormann (2005). One method is to take the mean-field limit to obtain an equivalent continuum model, where the positions and orientations of cells are translated into cell density and mean orientation across continuous space (Albi and Pareschi, 2013; Degond and Motsch, 2008; Großmann et al., 2016). This has been done for the Vicsek model (Bolley et al., 2012), but is mathematically challenging and poses challenges for discontinuous coefficients, such as those that arise in Leech et al. (2024). Instead of considering the large-cell-number limit, in this work we look in the other direction and consider two interacting cells, with the goal of making analytic progress. Since the agent-based model in Leech et al. (2024) requires computing the points of intersection between the boundaries of overlapping ellipses, it is helpful to impose some symmetry in relative ellipse orientation and position to facilitate analysis. Specifically, this allows us to analytically determine the overlap points of the two overlapping ellipses, which allows us to write down explicit governing equations. We are able to make significant analytic progress in understanding the non-trivial dynamic interaction between two interacting cells, and how the different aspects of the model contribute to alignment between two cells.

### Structure of this work

In this work, we will mathematically analyse the model derived in Leech et al. (2024) by considering two interacting cells with some symmetry imposed. We begin by introducing the full model, and then derive the analytical framework which leads to a coupled dynamical system for three time-dependent scalar quantities: distance between the cells, relative cell orientation and cell aspect ratio (Sec. 2). Analysis of this three dimensional (in variable space) dynamical system is done in Sec. 3. We then reduce the system to two dimensions (in variable space) by taking the limit of rigid-cell-shapes (i.e. fixed aspect ratio), which allows us to fully understand the effect of the self-propulsion speed on alignment, as well as to quantify the dependence of alignment strength on various model parameters (Sec. 4).

## 2 Model derivation

### 2.1 Full model summary

In Leech et al. (2024), an energy minimisation approach was used to derive a system of governing equations that describe the behaviour of a collective of self-propelled ellipse-shaped cells, moving in two spatial dimensions, that strive to avoid cell overlap upon interacting with one another. Full details of the derivation and equations for cell collectives can be found in Leech et al. (2024). Here we only summarise the most important aspects and provide equations for the interaction of two cells.

#### Description of the cells

In the following all quantities are non-dimensionalised with respect to a reference length 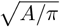 and reference time *Aη/σ*, where *A* is the cell area, *η* is the strength of friction that a cell experiences with the substrate, and *σ* is the strength of overlap avoidance. A cell is then characterised by its (non-dimensional) position **X** ∈ ℝ^2^, its orientation *α*, and its aspect ratio *r*, see Fig. 1A. Each material point inside the elliptic cell is described by the parameters *s* ∈ [0, 1] and *θ* ∈ [0, 2*π*) by

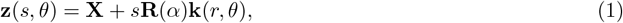

where the rotation matrix **R** and the shape vector **k** are given by

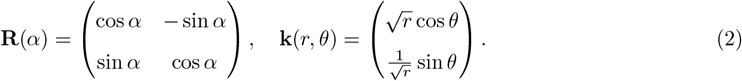

Note that this parametrisation leads to an area element *s d*(*s, θ*), which is independent of *θ* and *r*. This implies that changes in aspect ratio do not affect the (assumed) homogeneous density of material points inside the cell.

#### Model equations

To obtain equations that describe how cells change their position, orientation and aspect ratio (while keeping their area constant), we assume their movement minimises a potential, which models friction, overlap avoidance, relaxation to a preferred aspect ratio 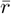 and self-propulsion. The governing equations for two interacting cells then are

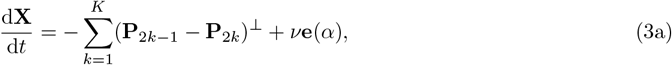

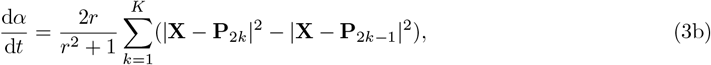

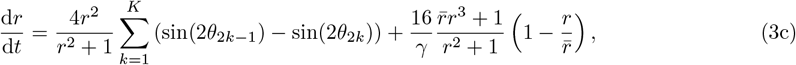

where **e**(*α*) = (cos *α*, sin *α*)^*T*^, ⊥ denotes the left-turned normal vector and

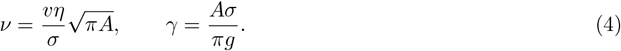

The non-dimensional quantities *ν* and *γ* defined in (4) depend on the self-propulsion speed *v*, the cell area *A*, the strength of overlap avoidance *σ*, the strength of friction with the substrate *η*, and the strength of relaxation to the preferred aspect ratio *g*. The quantity *ν* can be interpreted as the ratio of the self-propulsion speed to the strength of overlap avoidance in the presence of friction. A larger value of *ν* means a faster self-propulsion, or a smaller overlap avoidance strength. The quantity *γ* can be interpreted as the ratio between the strength of overlap avoidance and the strength of shape restoration. A larger value of *γ* means that the strength of relaxation to the preferred aspect ratio *g* is smaller and hence cell aspect ratios away from 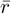 will be punished less. The pairs of points where the boundaries of the two cells intersect are given by (**P**_2*k*−1_, **P**_2*k*_), with *k* = 1, … *K. K* = 0, 1 or 2 indicates the number of intersection point pairs (the case of one or three intersection points can be reduced to the case of zero and two intersection points respectively). The ordering of intersection points is shown in Fig. 1B. The angles *θ*_*j*_ in (3c) correspond to the *θ* values that parameterise the intersection points **P**_*j*_, i.e. **P**_*j*_ = **z**(1, *θ*_*j*_) in (1).

#### Interpretation

For ease of interpretation we refer to the case of only one pair of intersection points, i.e. *K* = 1. This is the situation depicted in Fig. 1C-D. From equation (3a), we see that the centre of the cell, **X**, is being pushed perpendicular to the vector connecting the points of overlap **P**_1_ and **P**_2_. We can see from (3b) that the cell is rotated by an amount proportional to the difference in the square of the lengths of the lines connecting the cell centre and the intersection points, thus turning the cell in the direction from the shorter to the longer line, Fig. 1C. The first term in (3c) (compare Fig. 1D) leads to cells shortening if the cell overlap is near the ends of the cells and lengthening if the overlap is along the sides. This will happen at a faster rate for larger values of *r*. The final term on the right-hand side of (3c) acts to restore the cell’s aspect ratio to the preferred aspect ratio 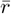. In this work we mostly focus on “long” cells 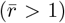, but will also consider “wide” cells 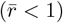 in the stability analysis of Sec. 3.

### 2.2 Symmetric cells - the analytical framework

To further analyse and understand system (3), we impose a symmetry condition on the two interacting cells. This allows us to obtain explicit expressions for the points of intersection, and therefore analytically tractable equations. We consider cell 1 with centre (*x*(*t*), *y*(*t*)), orientation *α*(*t*) and aspect ratio *r*(*t*) interacting with cell 2 with centre (*x*(*t*), −*y*(*t*)), orientation −*α*(*t*), and aspect ratio *r*(*t*) as shown in Fig. 2C. Both cells have equal self-propulsion parameters *ν*. The points belonging to the two cells are then parameterised as in (1) by

**Figure 2:**
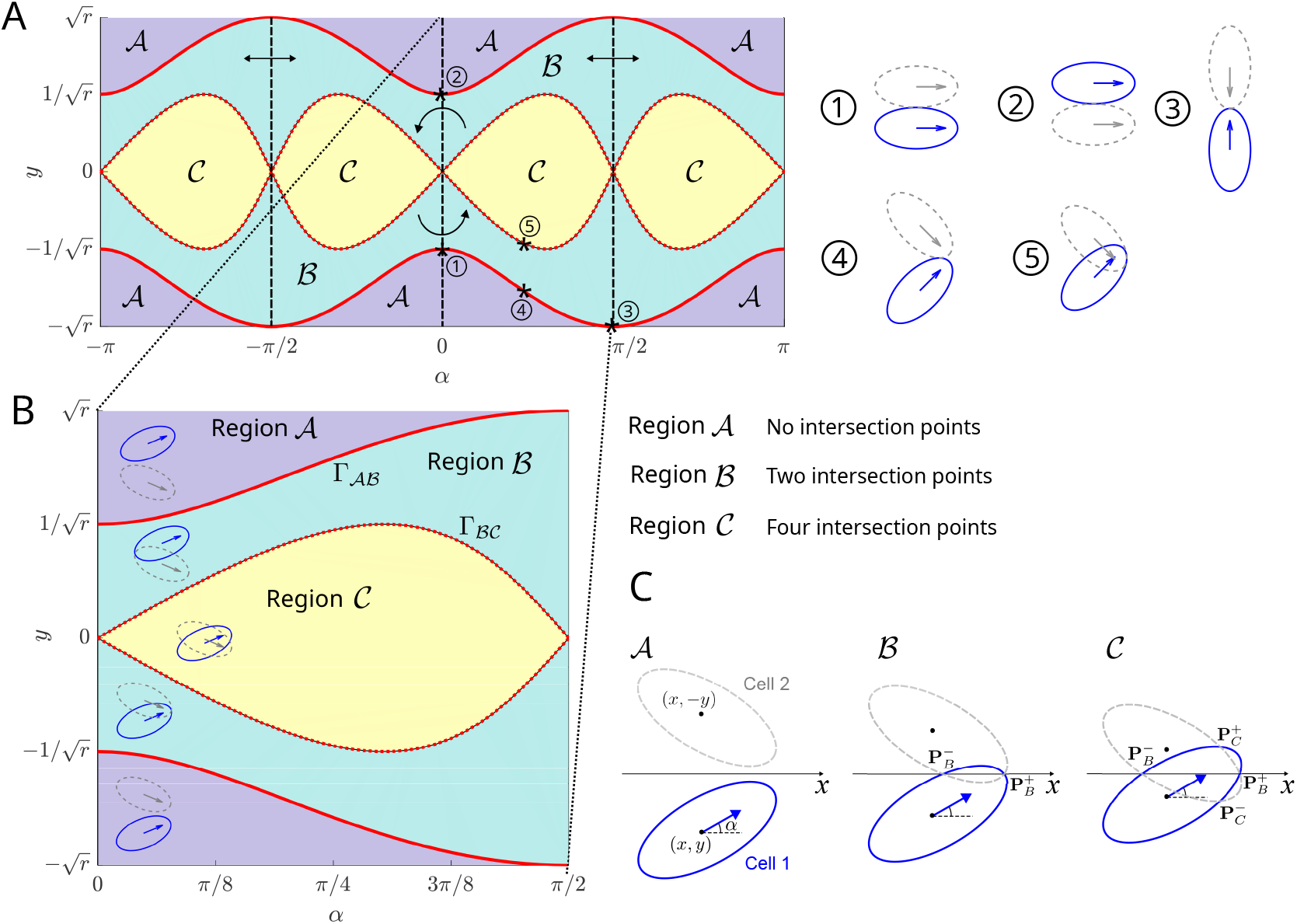
A: The (*α, y*)-plane for *r* = 2 with rotational and reflexive symmetries marked. Numbers 1-5 show example cell configurations on region boundaries. Regions 𝒜, ℬ and 𝒞 are coloured in purple, green and yellow respectively. B: Zoom into *α* ∈ [0, *π/*2] with regions 𝒜, ℬ and 𝒞 marked and typical cell configurations depicted. C: Notation for cell centres, orientation and intersection points.

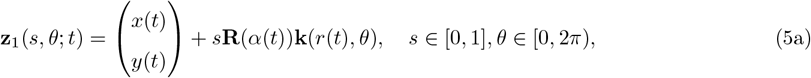

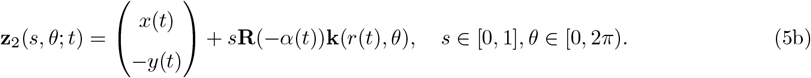

By considering where the cell boundaries *θ* ↦ **z**_1_(1, *θ*; *t*) and *θ* ↦ **z**_2_(1, *θ*; *t*) intersect (neglecting borderline cases), the two cells can have zero, two or four points of intersection. We obtain the following points of intersection (in Cartesian coordinates, relative to the cells’ *x*-position), see Fig. 2C:

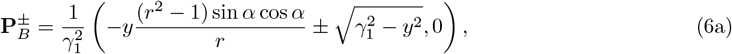

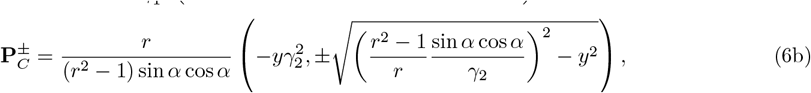

where we have defined

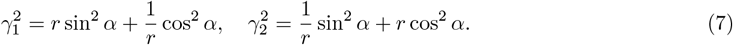

Given the square roots in (6a) and (6b), these points of intersection only exist in certain regions of (*α, y, r*)-space. We denote these regions by 𝒜, ℬ and 𝒞, see Fig. 2A,B: In region 𝒜 there are no points of intersection, in region ℬ there are two points of intersection, 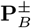, and in region 𝒞 there are four points of intersection, 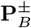 and 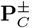. The boundaries between these regions occur when the square root terms in (6a) and (6b) vanish, which allows us to write these boundaries using the curves

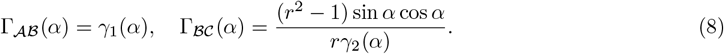

We can now formally define the regions via

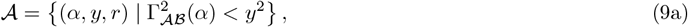

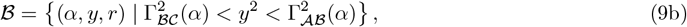

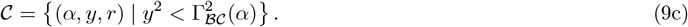

With (9) we can substitute the points of overlap in (6a) and (6b) into the two-cell versions of the full governing equations (3) to formulate the explicit two-cell governing equations for (*α, y, r*) ∈ ℝ × ℝ× ℝ^+^ with given initial conditions (*α*(0), *y*(0), *r*(0)) = (*α*_0_, *y*_0_, *r*_0_).

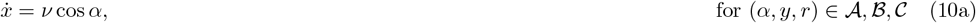

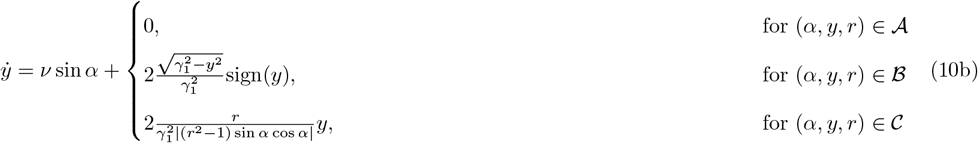

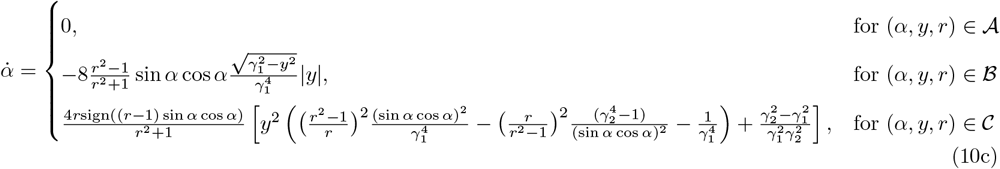

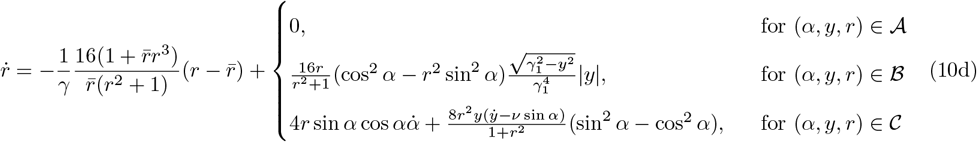

This is a rather complicated system of coupled non-linear differential equations, however the following sections will show that we can make significant (analytical) progress in understanding its behaviour. We start by discussing the general behaviour of the system (10) to gain initial insight, before presenting several analytical results in Secs. 3 and 4. We note that the system is invariant to the transformation (*α, y*) → (−*α*, −*y*), i.e. there is rotational symmetry about the line *α* = *y* = 0. We also note that the system has reflectional symmetry about *α* = *π/*2 and *α* = −*π/*2, see Fig. 2A. We can consequently restrict our analysis to the region *α* ∈ [0, *π/*2], see Fig. 2B, since the results can be extended to the full *α*-range via symmetry arguments. To aid interpretation of the dynamical system, we indicate various physical configurations of the cells in (*α, y*)-space in Fig. 2.

#### Interpretation of equations

With a better understanding of the phase space, and how this corresponds to cell configuration, we now discuss the equations in (10). Equation (10a) is decoupled from (10b), (10c) and (10d), hence it suffices to analyse the (*α, y, r*)-system. The first terms in (10a) and (10b) represent self-propulsion that leads to movement in the direction of (cos *α*, sin *α*). This is proportional to the non-dimensional parameter *ν* that has been defined in (4). Inspecting (10b) further, we note that if cell 1 is lower than cell 2 (*y <* 0), then the second term in (10b) is negative and cell 1 will move downwards, and vice versa for *y >* 0. This is a result of overlap avoidance pushing the cells apart. Equation (10c) describes how the cell orientation changes over time. If we consider *r >* 1, *α* ∈ (0, *π/*2), we see that 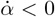 in region ℬ. This means that overlap avoidance in region ℬ causes a clockwise rotation and drives the system towards *α* = 0, i.e. towards velocity alignment. Conversely, for 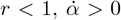 in region ℬ and the cells become less aligned. We will revisit this dependence of the behaviour on *r* when inspecting the stability of steady states, see also Fig. 5. In region 𝒞, both signs of 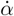 are possible. The first term of (10d) describes how the cells will relax back to their preferred aspect ratio 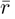, with 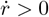 0 if 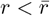 and 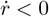 if 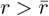 We see that in regions ℬ and 𝒞, overlap avoidance causes the aspect ratio *r* to change. If cos^2^ *α* − *r*^2^ sin^2^ *α >* 0 then 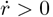. This will happen for example when the cells are side-by-side (*α* ≈ 0) and hence strive to increase their aspect ratio (for *r >* 1 this would mean elongation) to avoid overlap. If cos^2^ *α* − *r*^2^ sin^2^ *α <* 0 then 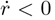, which would for example occur when cells are head-to-head (*α* ≈ *π/*2). In this case the cells will decrease their aspect ratio (for *r >* 1 this would correspond to shortening) to avoid overlap. The behaviour in region 𝒞 is more complex in general.

## 3 Deformable cells: *γ >* 0

We first explore the full shape-change model (10) where *γ >* 0 and *ν* = *O*(1). This involves explicitly accounting for the restoration time of the aspect ratio *r* to its preferred value 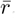, where *γ* can be thought of as the timescale of shape restoration. The full model is a 3D (in variable space) dynamical system where *y, α* and *r* vary in time in response to cell overlap, self-propulsion and shape restoration. Ignoring movement in the *x*-direction, system (10) has steady states at the points 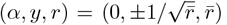. This corresponds to the cells having their preferred aspect ratio 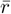 and being positioned side-by-side, with their orientations parallel and their boundaries just touching, illustrated in Fig. 2A, examples 1 and 2. We note that there are additional steady states, but only consider the two listed above in the subsequent analysis since these correspond to cell alignment, which is the focus of this work. To determine the stability of these steady states, we perturb the system around the points 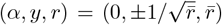. Importantly, a standard linear stability analysis would not be sufficient to determine their stability. In fact, such an analysis would be degenerate, and determining the stability of these states is non-trivial, as we will see below.

### 3.1 Stability analysis

We consider the steady point 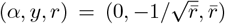 (Fig. 2A, example 1), noting that the analysis of the steady point 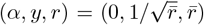 will follow via the rotational symmetry of the system. In order to perform the stability analysis, we perturb the steady point 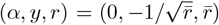 by a small amount (to be defined below) and calculate the subsequent dynamics of the system. This steady point is degenerate, so the scalings of our perturbation and its subsequent dynamics are non-standard.

#### Defining the perturbations

To capture the richest dynamics, we consider the distinguished asymptotic limit in which as many mechanisms as possible balance at the same time. If we define the (small) perturbation in *α* to be of *O*(*ε*), where *ε* ≪ 1, then with the benefit of hindsight and justified *a posteriori*, distinguished asymptotic limits occur when *γ* = *O*(*ε*) and when *γ* = *O*(1*/ε*). The former is the physically relevant case, since *γ* = *O*(1*/ε*) would allow for larger deformations to the aspect ratio *r*, which are not observed biologically. We henceforth focus on the distinguished limit *γ* = *O*(*ε*) and therefore scale *γ* = *ε*Γ, where Γ = *O*(1). In this case, the appropriate asymptotic scalings for the perturbations (justified *a posteriori*) are

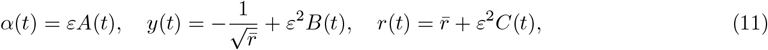

where *A, B*, and *C* are perturbations in their respective variables, and are functions of time that we will calculate. Understanding their dynamical behaviours will determine the stability of the steady point. To get an idea of the asymptotic structure of the solution before we go into the details, it is helpful to note that there are two distinguished timescales of interest in the system: the ‘early time’ where *t* = *O*(*ε*) and the ‘late time’ where *t* = *O*(1*/ε*). Over the early timescale, we will show below that *A*(*t*) remains unchanged, *B*(*t*) is driven by overlap avoidance and *C*(*t*) is driven by overlap avoidance and restoration to aspect ratio 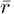. Using the early time results, we will then show that over the late timescale, *A*(*t*) is affected by overlap avoidance and, along with *B*(*t*) and *C*(*t*), decays to zero algebraically, demonstrating that the system is stable.

#### The dynamics of the perturbations

We now substitute (11) into (10) and note that since the steady state lies at the boundary of region 𝒜 and region ℬ, the perturbations could push the system in either of those two regions. We obtain

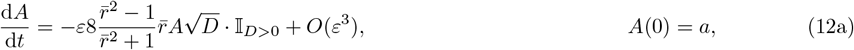

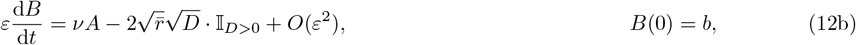

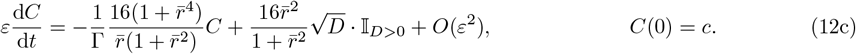

where 𝕀_*D>*0_ = 1 for *D >* 0 and zero otherwise, and *D* defined as

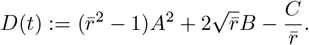

The sign of *D* determines whether we are in region 𝒜 or region ℬ and hence whether overlap avoidance takes effect. We now analyse the system (12), starting with the early time.

#### Early time

We start our analysis under the early timescale *τ* = *O*(1), defined via *t* = *ετ*. We indicate the early timescale variables with overhats, and therefore write

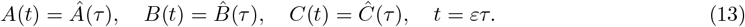

On substituting (13) into our governing equations (10) and taking the limit *ε* → 0, we obtain the following leading-order equations.

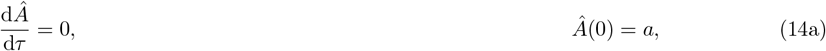

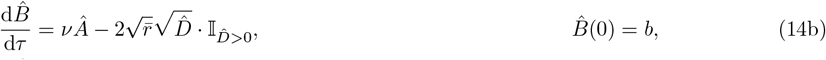

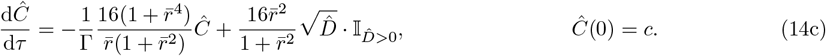

where 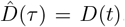. It is straightforward to use (14a) to determine that *Â*(*τ*) = *a* over the early time, and hence to deduce that the orientation is not affected over this timescale. The remaining system (14b)– (14c) governs the dynamics of 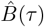 and 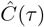, and can be solved computationally. The first term on the right-hand side of (14b) indicates that self-propulsion causes a change in 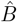 over the early time depending on the sign of *a*. This will be an increase if *a >* 0 since this means that the cell is inclined slightly upwards.

The second term on the right-hand side of (14b) represents overlap avoidance, suggesting that cells will move apart to avoid overlap, consequently causing a decrease in 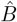. The relative size of these two terms will determine whether 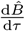 is initially positive or negative. The first term on the right-hand side of (14c) leads to a decrease in magnitude of 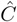, essentially restoring *r* to 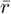. The second term on the right-hand side of (14c) is positive, corresponding to an increase in aspect ratio to avoid overlap. This forcing occurs because the cells are close to a side-by-side configuration near the steady state, and therefore increasing the aspect ratio reduces overlap. We also note that the non-trivial dynamics over this early timescale justify the *ε* scalings we initially imposed in (11).

#### Early-time overlap dynamics

To determine in which region, 𝒜 or ℬ the early time solution lies, it is useful to inspect the change in time of 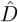, since its sign determines whether there is overlap 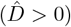 or not 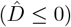. We obtain

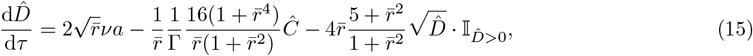

which, together with (14c) forms a nonlinear system of two autonomous ODEs. Phase plane analysis reveals that there is a qualitative difference for *a >* 0 and *a <* 0, see Fig. 3. We see that for *a >* 0, i.e. a slightly upward inclined cell 1, the dynamics will lead to 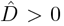, i.e. we end up in region ℬ and the two cells will eventually interact, even if they initially did not. Interestingly it is possible that cells initially interact, then stop interacting for a short time, and then interact again. Such intermediate short non-interaction periods can be caused by the relaxation to the preferred aspect ratio, see example trajectory in Fig. 3A. If on the other hand *a <* 0, then 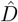 will eventually become negative and cells will stop interacting, irrespective of whether they did so initially. We additionally note that for *a <* 0 it is possible that cells that did not interact initially, then interact for a short amount of time due to shape relaxation, before ceasing to interact again, see example trajectory in Fig. 3B.

**Figure 3:**
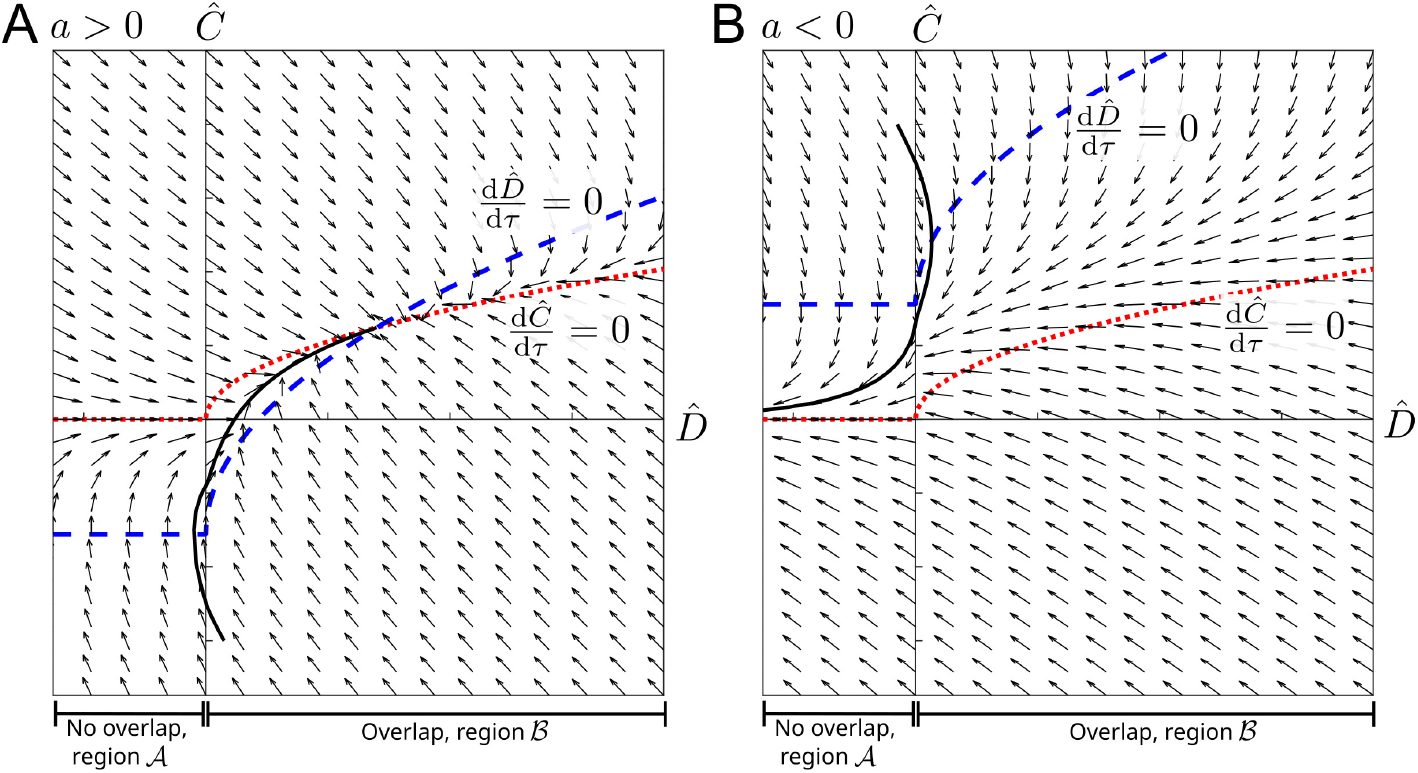
Early time dynamics: Phase portrait for the 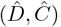-system (14c), (15) for *a >* 0 (A) and *a <* 0 (B). Nullclines are marked in dashed-blue 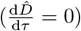 and dotted-red 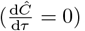. An example solution trajectory is shown in solid-black. 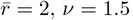.

#### Early-time limiting behaviour

For *a <* 0, cells will eventually stop to interact and move apart, hence the steady state is unstable for such perturbations, compare Fig. 5B, example 2. If *a >* 0, system (14) becomes independent of early time *τ* in the large-*τ* limit. We can calculate the specific constants to which the solutions tend by setting the left-hand sides of (14b)–(14c) to zero and solving the resulting algebraic equations. This procedure yields the following *τ* → ∞ results

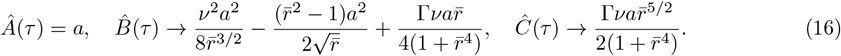

The early time ‘far-field’ conditions (16) will be required to asymptotically match into the late-time dynamics we consider next. Note that for these limiting values, the cells are still in region ℬ.

#### 3.1.2 Late time

Since we have already shown instability for *a <* 0, we only consider *a >* 0 here. Further, based on the discussion above we can assume that solutions are in overlap region ℬ. System (12) behaves as the early time far-field (16) until *t* = *O*(1*/ε*), when we have a new distinguished timescale that we refer to as the ‘late time’. By inspecting (12), we note that this is the timescale over which the dynamics of *A* become relevant. Formally, the late timescale is defined by the new variable *T* = *O*(1), where *t* = *T/ε*. We retain the perturbation scalings (11), but now use tildes to denote late-time variables, defining

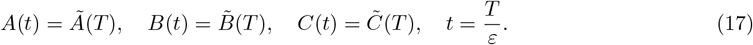

Over this timescale, in the limit *ε* → 0, our governing equations (12) become

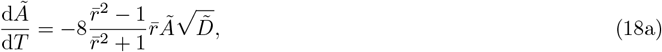

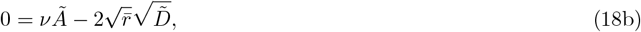

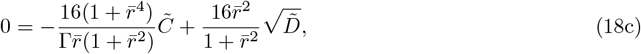

where

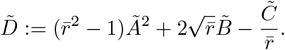

The ‘initial’ conditions are obtained by matching with the early-time far-field conditions (16),

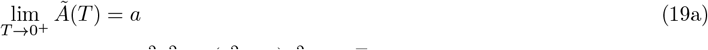

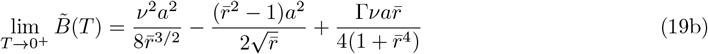

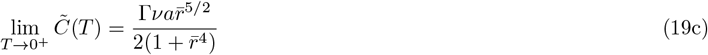

Although the differential-algebraic system (18)–(19) is nonlinear, we can simplify the nonlinearity in the differential equation (18a) using the algebraic equation (18b). This procedure reduces (18a) to the following nonlinear but separable differential equation in *Ã*

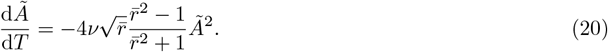

Solving (20), and substituting into the remaining algebraic equations (18b)–(18c), we obtain the late-time solutions:

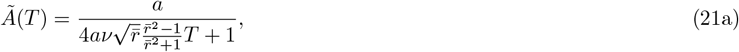

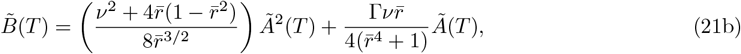

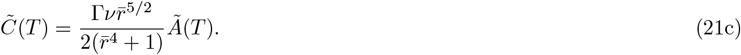

For 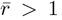, (21) shows that 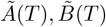 and 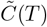 all decay to zero algebraically as *T* → ∞, compare Fig. 5A,B, example 1. The factor in front of *T* in (21a) is an increasing function of 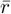 (for 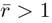), suggesting that larger preferred aspect ratios will lead to faster decay and hence faster alignment. If on the other hand 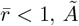 will blow up in finite time, indicating an unstable situation, compare Fig. 5C,D.

#### 3.1.3 Composite solution

Combining the solutions (14) and (21) in both timescales, we can obtain a uniformly regular (additive) composite asymptotic solution (Van Dyke, 1975) for *y, α* and *r* in terms of *t*.

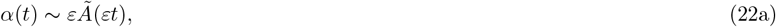

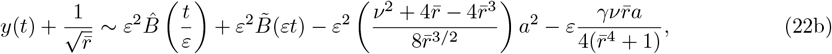

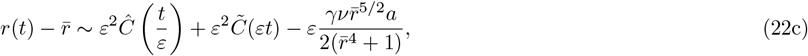

where the hatted (early-time) variables are defined in (14) and the tilded (late-time) variables are defined in (21). In Fig. 4 we verify that the quantities *A, B* and *C* all decay to zero as *t* → ∞ for *a >* 0 and 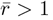, demonstrating that the point 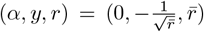 is stable in this case. On comparing our analytical solution for the stability analysis with the computational full solution, solved using ode15s in MATLAB, we see that we have a good agreement between the two for *ε* = 0.1 and for *γ* = 0.01, 0.1, 1. The fact that the approximation remains good also for larger values of *γ* is expected from our analysis since *γ* = *O*(*ε*) is a distinguished limit of the system. Hence we expect our analysis to hold in its sublimits until a new distinguished limit is reached. Specifically for this problem, our above analysis allows us to straightforwardly understand the subcases *γ* ≪ *ε* and *ε* ≪ *γ* ≪ 1*/ε* as regular sublimits of our analysis. We also note that the solution for *A*(*t*) is unaffected by *γ*, suggesting that having shape change in the model does not affect the change in orientation that two cells will experience in response to overlap avoidance. Since a change in orientation is needed for cells to align with each other, this suggests that varying the non-dimensional shape change parameter *γ*, at least if *γ* = *O*(*ε*) and *ν* = *O*(1), has little effect on cell alignment.

**Figure 4:**
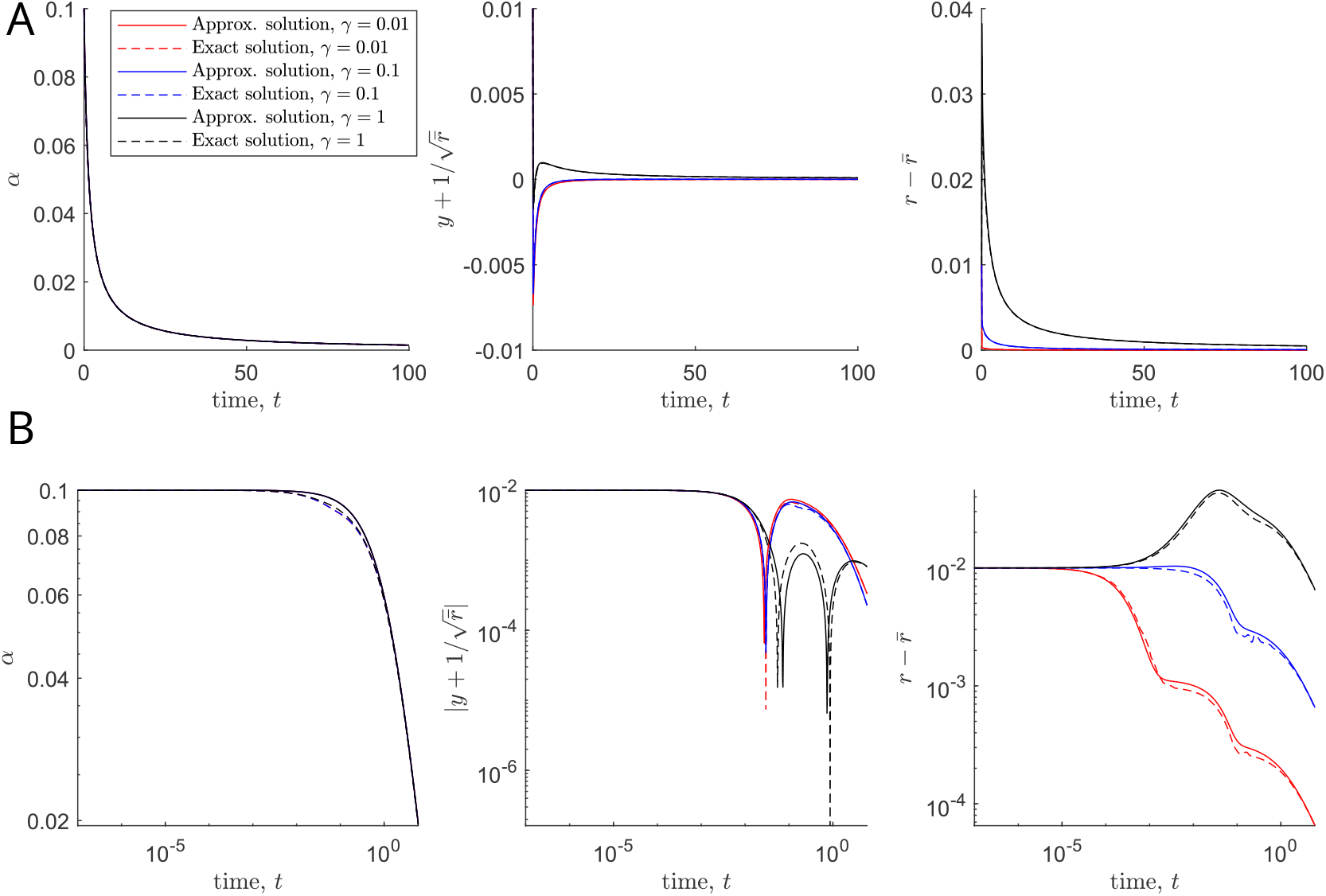
A, B: Plots of the exact solution of system (10) and the approximate solution given in (22) against time until *t* = 100 (A) and until *t* = 6 in a *log-log* plot (B), for *ε* = 0.1, *γ* = 0.01, 0.1, 1. Legend applies to all plots. Other parameters: 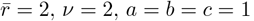

#### 3.1.4 Summary and discussion of stability results

Together we have shown that the steady state 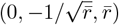 is half-stable for 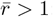 If the point is perturbed in the positive *α*-direction, then the perturbation will decay. If it is perturbed in the negative *α*-direction, perturbations will grow and cells will eventually stop interacting and move away from each other, see Fig. 5A,B. We also emphasize that the solutions (21) yield an algebraic decay for *a >* 0, rather than an exponential decay that might arise from a standard linear stability analysis. This *a posteriori* justifies our claim that a non-standard stability analysis is required to determine the system stability. For 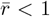 the point is always unstable: For *a <* 0 cells again eventually stop interacting and move away from each other. For *a >* 0, we can see that the orientation *α* moves away from away from *α* = 0 and increases. While we cannot use our approximate solutions to determine the limiting behaviour in this case, numerical results suggest that the solution converges to 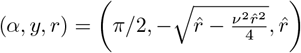 for some 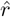, such that the limit is another steady state of (10b)-(10c). In this situation the cells face each other and self-propulsion, overlap avoidance and shape relaxation balance such that their distance stays constant, see Fig. 5C,D. Since this steady state is less biologically relevant, we do not systematically investigate its stability, but we expect it to be stable for 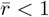. Using the rotational symmetry of our phase space, we obtain that the other steady state 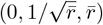 is also half stable for 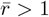 and unstable for 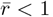.

**Figure 5:**
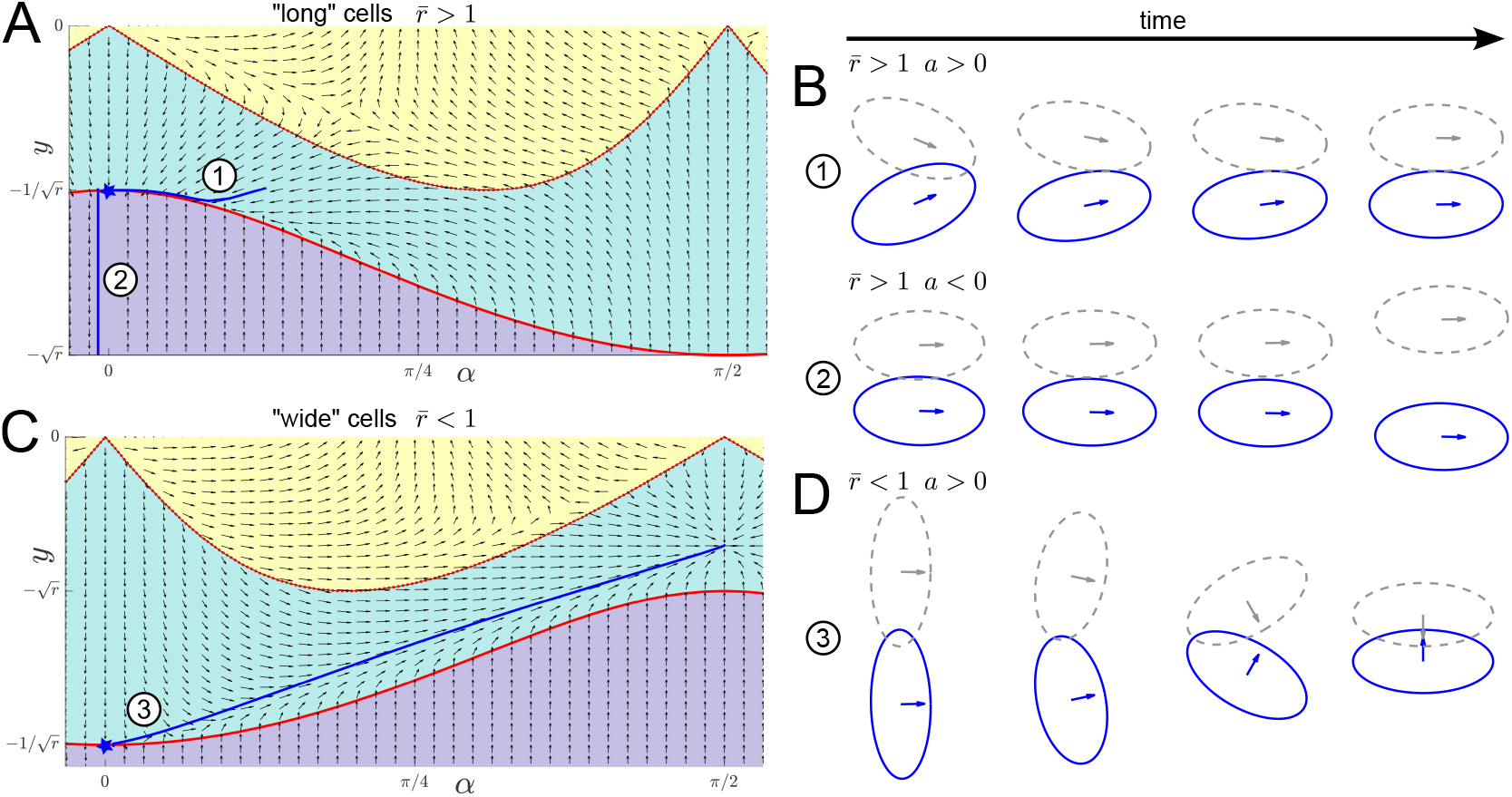
A, C: Typical phase portrait (omitting the *r*-direction) in (*α, y*)-space for long cells, 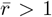, (A) and wide cells, 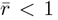, (B) with the analysed steady state marked with a blue star. Example trajectories (omitting the *r*-component) are shown in blue, numbers correspond to plots in B and D. B,D: Cell shapes at various time points corresponding to the trajectories marked in A and C. Other parameters: 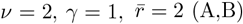 and 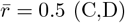.

### 3.2 Asymptotic sublimits

From the above stability analysis, we can understand why *γ* = *O*(*ε*) is a distinguished asymptotic limit of the system (i.e. a ‘least degenerate’ limit where as many processes as possible occur over the same timescale). Specifically, the natural overlap avoidance timescale is *t* = *O*(*ε*), and the aspect ratio restoration timescale is *t* = *O*(*γ*). The distinguished limit therefore arises because the timescales of these different processes coincide when *γ* = *O*(*ε*). Since there is an additional natural timescale in the system of orientation response when *t* = *O*(1*/ε*) (the ‘late’ time in our analysis above), we can deduce that there is a different (additional) distinguished limit when *γ* = *O*(1*/ε*). This additional distinguished limit would correspond to extremely “squishy” cells where shape change is punished very little, generating cells with aspect ratios far away from the natural shape 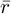 as a result of overlap avoidance. As this is less biologically relevant, we do not consider this case further.

Since distinguished limits correspond to maximal interaction of the natural processes, our asymptotic results for *γ* = *O*(*ε*) above will also hold for *γ* ≪ *ε* and *γ* ≫*ε* (i.e. ‘sublimits’ of the distinguished asymptotic limit), until new distinguished limits of *γ* are reached. It is instructive to briefly summarize what happens in the sublimits of the distinguished limit *γ* = *O*(*ε*) we analysed above.

#### Sublimit

*ε* ≪ *γ* ≪ 1*/ε*. In the sublimit of *ε* ≪ *γ* ≪ 1*/ε*, equivalently 1 ≪ Γ ≪ 1*/ε*^2^ (the upper limits of which correspond to the different distinguished limit noted above), the different balancing mechanisms over the early timescale diverge into two separate timescales: *t* = *O*(*ε*) and *t* = *O*(*γ*). Over the standard early timescale *t* = *O*(*ε*), only overlap avoidance affects *C*. Over this timescale, *B* and *C* grow without a restoring force. The timescale over which this restoring force becomes important is then a new intermediate timescale *t* = *O*(*γ*), which kicks in when *C* gets large enough for the overlap avoidance term to balance the aspect-ratio-restoring term. Over this intermediate timescale, *B* and *C* reach the equivalent steady states to those in the full behaviour we determined above. Finally, the late timescale *t* = *O*(1*/ε*) is essentially unchanged from the full analysis in this sublimit; it is this timescale over which *A*(*t*) varies. The full analysis for *γ* = *O*(1) is outlined in Appendix A.1.

#### Sublimit

*γ* ≪ *ε*. The opposite sublimit of *γ* ≪ *ε* (equivalently Γ ≪ 1), corresponds to cells that restore their natural aspect ratio very quickly (over a timescale of *t* = *O*(*γ*)), and then behave as non-deformable cells with distinct early (*t* = *O*(*ε*)) and late (*t* = *O*(1*/ε*)) time behaviours. This sublimit is tractable for a more general dynamic analysis, not just near the steady points, and is of biological interest. Therefore, we examine this case in more detail in Sec. 4.

### 3.3 Lessons for (collective) cell alignment

#### Alignment stability of “long” cells

The points 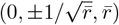 represent perfect alignment of two cells. For 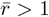 (“long” cells), the fact that these points are half-stable shows that alignment is sensitive to some perturbations, and suggests that overlap avoidance drives alignment, but that perfect alignment is fleeting in practice. In the many-cell version of the model in Leech et al. (2024), interactions with a third cell could perturb the perfect alignment between two cells in the unstable direction, i.e. such that they point away from each other (corresponding to *a <* 0 in the above analysis), in which case they would cease to interact and alignment would be broken. This could also occur if cell orientation is subject to noise, as would be the case in many realistic biological systems. This could explain why we see pockets of alignment, but no global alignment in the simulations in Leech et al. (2024).

#### The role of the aspect ratio and deformability

We found that for 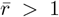 the decay to perfect alignment is faster for larger 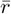, indicating that larger preferred aspect ratios lead to faster alignment. We will revisit the question of how the strength of alignment depends on the aspect ratio in Sec. 4. The stability analysis also showed that, near the steady state, the change in orientation is independent of the shape change parameter, suggesting that deformability does not aid alignment. The latter result is in contrast to the collective dynamics results of Leech et al. (2024), where deformability lead to more collective alignment. This indicates that the increased collective alignment due to deformability might be a consequence of the many-cell system, not of the alignment mechanism itself. The case 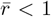 describes “wide” cells that move orthogonal to their long axis, such as keratocytes (fish fibroblasts). Here the perfect-alignment steady state is always unstable. This could indicate that we do not expect to see velocity alignment in such cells. However, it is also possible that the imposed symmetry condition in this work is not appropriate in this case.

#### Non-deformable cells, *γ* = 0, *r >* 1

In this section, we consider the rigid-cell-shape limit, taking *γ* → 0. Further, based on the findings in Sec. 3 we focus on “long” cells and hence assume 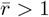 throughout this section. As noted in the previous section, there is a very fast timescale of *t* = *O*(*γ*) over which the cells restore and then maintain their preferred aspect ratio of 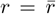. We will proceed by taking *γ* = 0 and 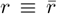, essentially analysing (10) after the very fast *t* = *O*(*γ*) timescale. This reduces (10) to a 2D (in variable space) dynamical system in *α* and *y*, (10b),(10c). This reduction is both biologically relevant and more straightforward to analyse and interpret. In the rest of this section, we will denote 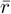 by *r* for notational convenience, where *r* no longer has a time dependence. The (*α, y*) state space is split into regions 𝒜, ℬ and 𝒞, see Fig. 2B. We start our investigation by confirming that we obtain the same stability result from Sec. 3 before exploring the resulting 2D dynamical system computationally. Then, to gain deeper insight into the precise effect of self-propulsion, which has dimensionless strength *ν*, we consider the case of *ν* = 0 before using asymptotic methods to understand the case where *ν* ≪ 1, noting that the limit *ν* → 0 is singular.

### 4.1 General behaviour

#### Stability analysis

For *ν >* 0, *ν* = *O*(1), the stability of 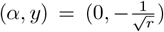 is inherited directly as a sublimit Γ → 0 of the full system stability analysis we conducted for deformable cells in Sec. 3 above. We assume the initial conditions are such that we stay in region ℬ (see discussion in Sec. 3). Specifically for *a >* 0, and if we take *γ* → 0 in (10d), or equivalently Γ → 0 in (14c), we find that *C*(*t*) → 0 over *t* = *O*(*γ*), which is much quicker compared to the other two timescales of *O*(*ε*) and *O*(1*/ε*). Then, over the fast timescale *t* = *O*(*ε*), (14) reduces to

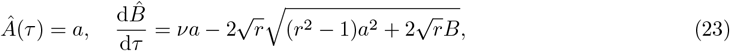

and over the slow timescale *t* = *O*(1*/ε*), (18) becomes

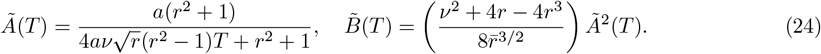

We can see that since 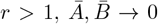 algebraically as *T* → ∞. Since perturbations with *a <* 0 will not decay, we therefore showed that, as for the system for deformable cells, the point 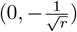 is half-stable.

#### Phase portrait

Next we explore the system more generally by computationally generating a phase portrait with some example trajectories in Fig. 6. Nullclines corresponding to 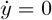 (pink) and 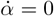 (green) are shown as dotted lines. We see that in region 𝒜 (purple) the arrows all point vertically upwards. This is because this is the region of no overlap and hence the cells will only change their position (as a result of self-propulsion) and not their orientation. The arrows are pointed upwards since *α* ∈ (0, *π/*2) and hence this leads to an increase in *y* since the cell is inclined upwards. If we focus our attention on the point 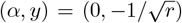, we see that all arrows in the surrounding area are directed towards this point, illustrating that this point is an attractor from the side where *α >* 0. Example trajectories are shown in blue and these indicate that there are two outcomes depending on the initial conditions: (1) the trajectory moves towards the half-stable point 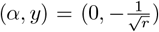, or (2) the trajectory crosses region 𝒞 and ends up leaving the overlap region with *y >* 0. This is a situation where cells crawl over each other, then stop interacting and continue to move away from each other. The line separating these two outcomes is a non-trivial separatrix, indicated by the solid blue line in Fig. 6B. We can calculate this separatrix computationally, by starting near the separatrix and solving the system (10) backwards in time (Fay and Joubert, 2010).

**Figure 6:**
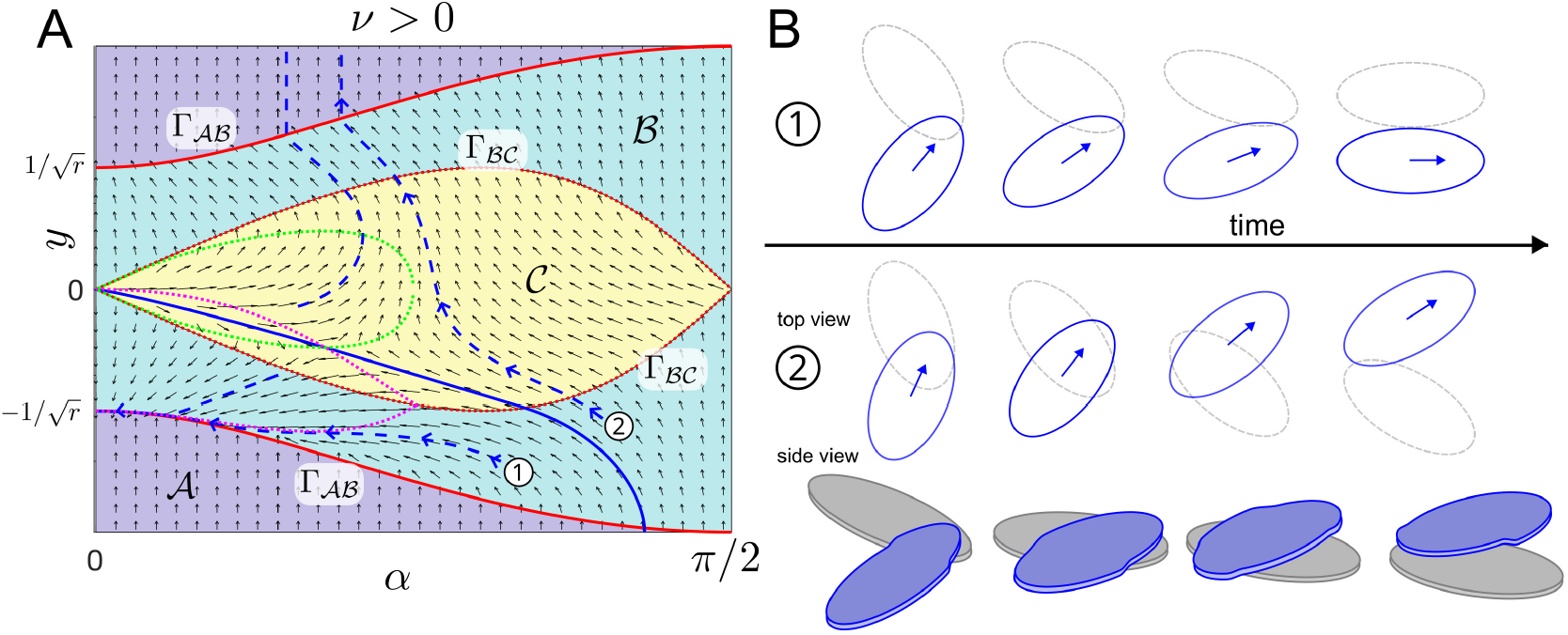
A: Phase portrait for *ν* = 2, *r* = 2 showing nullclines (dotted pink and green), boundaries between regions (red), the separatrix (solid-blue) and example trajectories (dashed-blue). B: Cell shape snapshots of the system over time for two example trajectories labelled (1) and (2) in A with a 3D side view included for (2). Note that which cell is above and which is below is arbitrary. The cell in blue corresponds to the trajectory in blue.

#### Quantification of alignment

Next, we want to quantify how favourable different sets of model parameters are for alignment. To do this we test initial conditions with 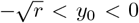 and 0 *< α*_0_ *< π/*2 (i.e. cell 1 underneath cell 2 and cells moving towards each other) and record the resulting post-interaction angle *α*_final_(*α*_0_, *y*_0_). This will either be the angle obtained once the interaction has ended after finite time, or the limiting angle for *t* → ∞, in the case of infinitely long interactions. Since *α*_final_(*α*_0_, *y*_0_) will lie between 0 and *π/*2, and 0 corresponds to perfect velocity alignment, we define the interaction strength for a particular parameter set by

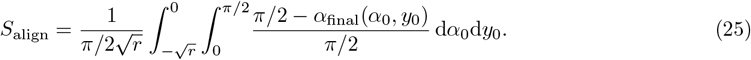

The alignment strength will be between 0 and 1, with *S*_align_ = 1 corresponding to perfect alignment for all used initial conditions. We determined *S*_align_ computationally by discretising the integral. Fig. 7A shows a non-monotonous dependence of alignment strength on the self-propulsion speed *ν*: For non-zero speeds, larger speeds lead to less alignment. Fig. 7E,F suggest this is because for larger *ν* fewer initial conditions lead to cells getting trapped in the region underneath the separatrix, where the interaction leads to perfect alignment. This is because larger speeds will more often allow cells to leave the *y <* 0 region, where they will stop interacting in finite time. However *ν* = 0 is not favourable for alignment. We will discuss this case in Sec. 4.2. The dependence of alignment strength on the aspect ratio *r* is also non-monotonic. The general picture is that both too round and too elongated cells shapes lead to less alignment. This can be understood by comparing Figs. 7C,F. On the one hand, larger aspect ratios lead to fewer initial conditions being in the perfect-alignment region underneath the separatrix. On the other hand, larger aspect ratios cause more post-interaction alignment (smaller values of *α*_final_(*α*_0_, *y*_0_)) for cells outside the perfect-alignment region.

**Figure 7:**
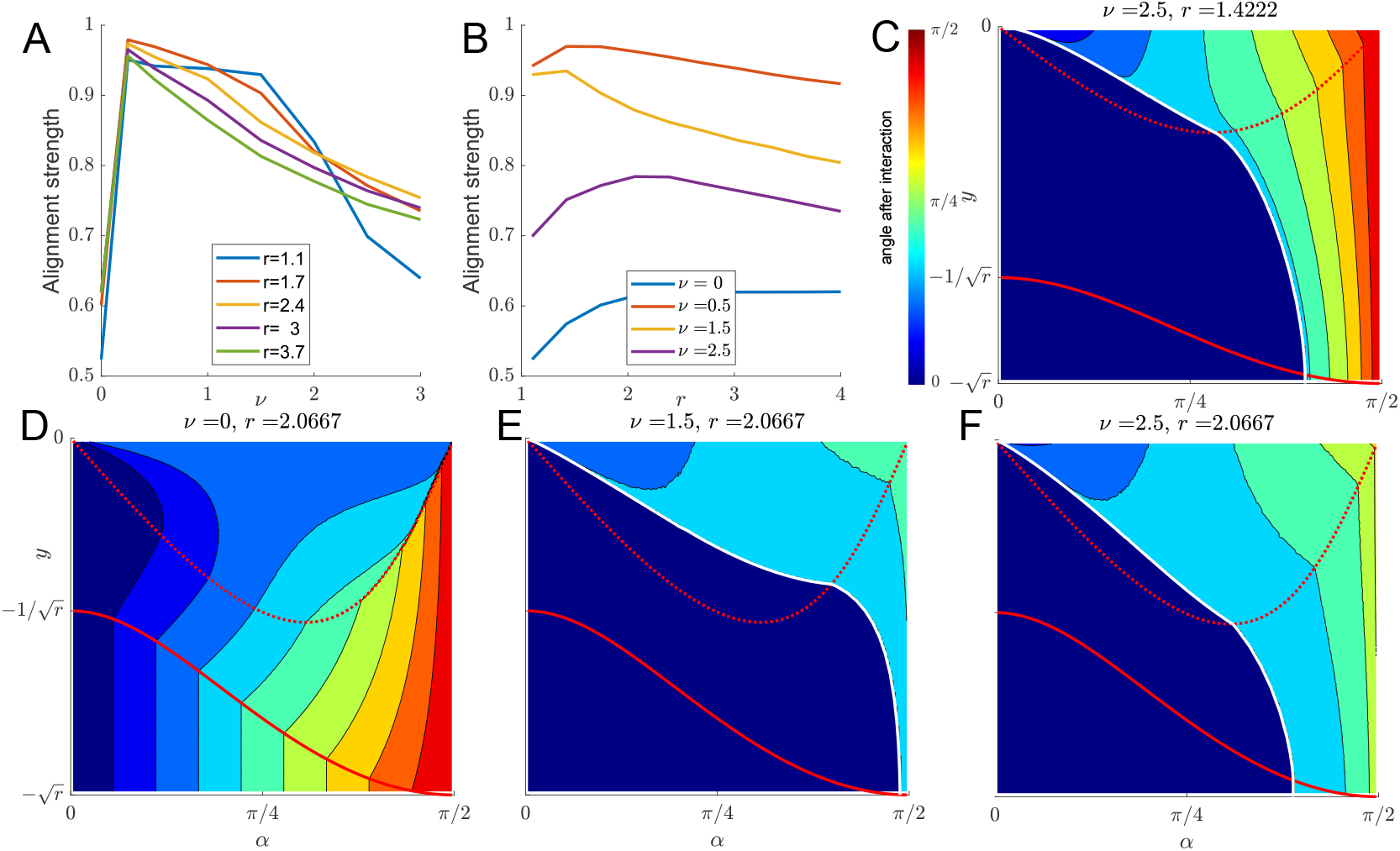
A-B: Alignment strength as defined in (25) as functions of *ν* (A) and *r* (B). C-F: Examples of interaction outcomes for different values of *ν* and *r*. Colour shows angle *α* after interaction has ended for the initial conditions at that position. Region boundaries are shown in red, the separatrix in white.

### 4.2 No self-propulsion, *ν* = 0

To investigate the effect of propulsion on (10) with *γ* = 0, we now analyse the singular change in the system in the limit of small *ν*. Specifically, we are interested in the strong difference in system behaviour for zero self-propulsion (*ν* = 0) and small self-propulsion (0 *< ν* ≪ 1) suggested by Fig. 7A,D. We start by considering zero self-propulsion, *ν* = 0. The governing equations (10) with *ν* = 0 are

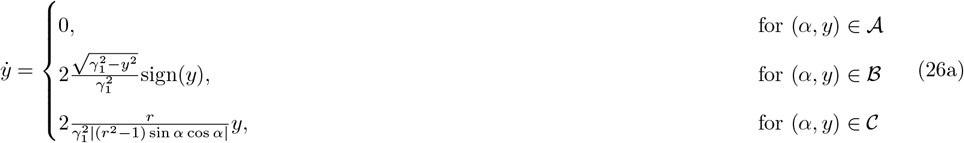

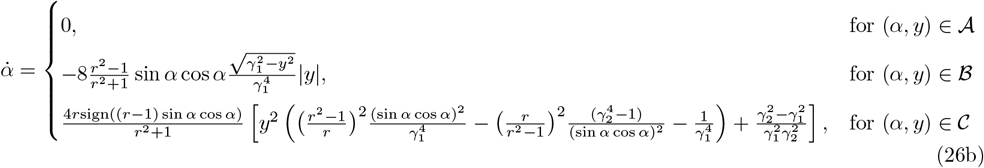

#### Summary of system behaviour

System (26) is symmetric around *y* = 0. Together with the symmetry considerations for *α*, it therefore suffices to consider *y* ≤ 0, *α* ∈ [0, *π/*2]. We inspect the corresponding phase portrait in Fig. 8. We find that 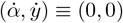 in region 𝒜, since in the absence of self-propulsion and shape relaxation, overlap avoidance is the only driver of change/movement. Throughout region ℬ and 𝒞 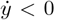, indicating cells move apart as long as they interact. Further the boundary between region 𝒜 and ℬ, given by *y* = −Γ_𝒜ℬ_ = −*γ*_1_ is now a line of stationary points. Together, this suggests the following solution behaviour, which we will prove in the next paragraph. If we let (*α*(0), *y*(0)) = (*α*_0_, *y*_0_) be the initial conditions and assume that *y*_0_ *<* 0, 0 *< α*_0_ *< π/*2, we find that:

**Figure 8:**
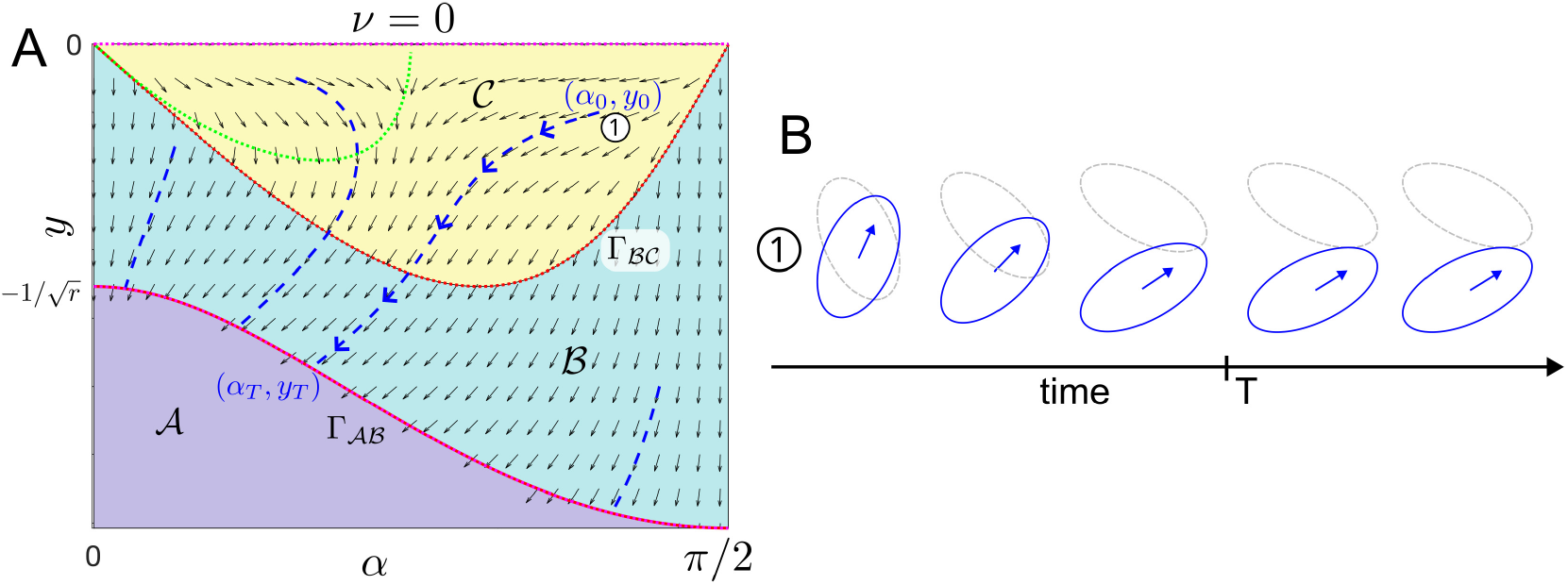
A: Phase portrait in (*α, y*)-space for *ν* = 0, *r* = 2. Example trajectories shown in blue. Boundaries between regions shown in red. Nullclines shown in green. B: Snapshots of the cell configurations over time corresponding to the trajectory in A marked with (1) at initial conditions (*α*_0_, *y*_0_).

i. If (*α*_0_, *y*_0_) lies in region 𝒜 (no interaction), the solution will remain at (*α*_0_, *y*_0_) for all time.
ii. If (*α*_0_, *y*_0_) lies in region 𝒞 (four intersection points), it will move into region ℬ in finite time and remain in region ℬ.
iii. If (*α*_0_, *y*_0_) lies in region ℬ (two intersection points) it will converge to a point (*α*_*T*_, *y*_*T*_) on the 𝒜-ℬ boundary in finite time, see trajectory in Fig. 8A,B.

#### Proof of system behaviour

Part (i) follows trivially from the dynamical system (26). Part (ii): From (26a) we see that 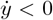 when *y <* 0 within region 𝒞 and on the boundary of region 𝒞. We also have that 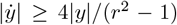 which shows that 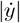 is bounded away from zero (note that *r >* 1). Hence, we can conclude that if (*α*_0_, *y*_0_) ∈ 𝒞 then there exists a finite time, at which the trajectory (*α*(*t*), *y*(*t*)) will cross the ℬ-𝒞 boundary and cross into region ℬ. Next we claim that region ℬ is invariant, i.e. that once a trajectory is in region ℬ, it remains in region ℬ. To show that this is the case, we characterise the solution curves in region ℬ. We consider the case where (*α*_0_, *y*_0_) and (*α, y*) are in ℬ and derive an equation for *y*(*α*). Dividing (26b) by (26a) we obtain

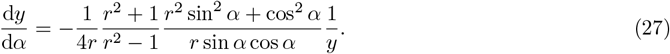

Equation (27) can be solved explicitly to yield

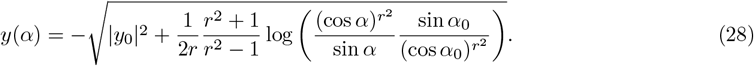

To show that region ℬ is invariant, we will show that once the trajectory moves from region 𝒞 into region ℬ, it is not possible for the trajectory to move back into region 𝒞 again. To do this, we will show that if we start on the boundary of region ℬ and 𝒞, given by 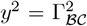 as defined in (8), we will enter region ℬ and will not intersect the line 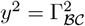 again, showing that it is not possible for the trajectory to cross this line a second time and enter back into region 𝒞. We substitute our initial conditions 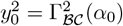 into (28) and want to solve 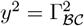 to see whether the trajectory *y* could intersect the boundary Γ_ℬ𝒞_ again. If there are no additional solutions to this equation, then the trajectory does not cross the boundary between regions ℬ and 𝒞 again and hence does not re-enter region 𝒞. From this procedure we obtain

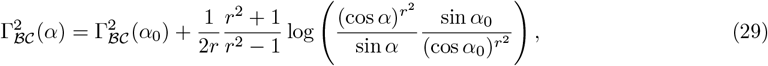

where Γ_ℬ𝒞_ is defined in (8). Rearranging (29) and defining *s* = sin *α*, we obtain

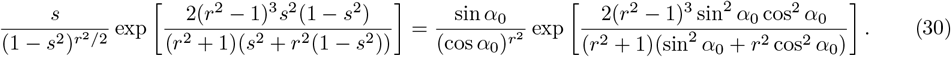

We define the left-hand side of (30) as *s* ↦ *f* (*s*) with domain [0, 1). Differentiating *f* with respect to *s* shows that *f* is a strictly monotonically increasing function of *s* with *f* (0) = 0 and *f* (*s*) → ∞ for *s* → 1^−^. Since the right-hand side is always positive, there exists a unique solution *s*_*sol*_ ∈ [0, 1) for each (*α*_0_, *y*_0_) and consequently a solution *α*_*sol*_ = arcsin *s*_*sol*_ ∈ [0, *π/*2] such that *y*(*α*_*sol*_) = −Γ_ℬ𝒞_(*α*_*sol*_). However, since we know that the initial condition satisfies these requirements, the unique solution is the initial condition i.e. *α*_*sol*_ = *α*_0_. Therefore, after intersecting the curve 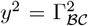 and moving into region ℬ, the trajectory will not intersect this boundary curve again. Hence, once the trajectory is in region ℬ it will remain in region ℬ.

For part (iii), we start by noting that the curve *y* = −Γ_*AB*_ describes a line of stationary points (where 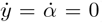). Showing that *y*(*α*) always intersects −Γ_𝒜ℬ_(*α*) is equivalent to showing that 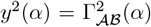 has a solution. Using *y*(*α*) as defined in (28) and Γ_𝒜ℬ_ as defined in (8) this can be written as

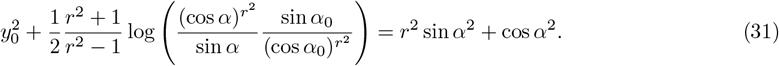

We need to show that (31) has a solution *α* ∈ [0, *π/*2]. We define *s* = sin *α* and use this to rewrite (31) as

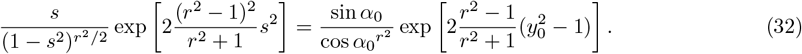

We define the left-hand side as *s* ↦ *h*(*s*) with domain [0, 1). Differentiating *h*(*s*) we find that *h*^*t*^(*s*) *>* 0 and can therefore conclude that *h* is a strictly monotonically increasing function of *s* with *h*(0) = 0 and *h*(*s*) → ∞ for *s* → 1^−^, i.e. its range is [0, ∞). Since the right-hand side is always positive, there exists a unique solution *s*_*T*_ ∈ [0, 1) for each (*α*_0_, *y*_0_) and consequently a solution *α*_*T*_ = arcsin *s*_*T*_ ∈ [0, *π/*2] such that *y*(*α*_*T*_) = −Γ_𝒜ℬ_(*α*_*T*_). Finally, we need to show that (*α*_*T*_, *y*_*T*_) is reached in finite time, where we define *y*_*T*_ := *y*(*α*_*T*_). To show this, we substitute (28) into (26b), the governing equation for 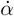, separate variables and integrate over time from 0 to *t*, obtaining

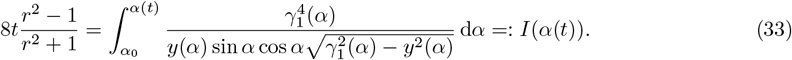

Our final task is to show that 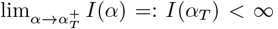, which will imply that (*α*_*T*_, *y*_*T*_) is reached in finite time 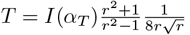. We use the substitution *s* = sin *α* in (33) and obtain

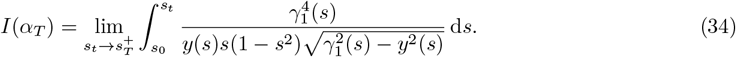

To understand whether (34) converges, we need to understand the nature of the singularity at *s* = *s*_*T*_ where 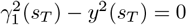. Taylor expanding this term around *s* = *s*_*T*_ gives

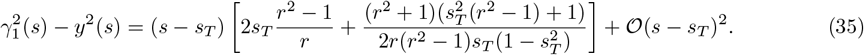

Since the coefficient of the linear *s* − *s*_*T*_ term in (35) does not vanish, the singularity in the integrand is an inverse square root, and therefore is integrable. That is, *I*(*α*_*T*_) *<* ∞ and hence (*α*(*t*), *y*(*t*)) = (*α*_*T*_, *y*_*T*_) for *t* = *T <* ∞. This explains the computational results shown in Fig. 7A,D, i.e. that for *ν* = 0 perfect alignment is typically not achieved.

### 4.3 Slow-moving cells, 0 *< ν* ≪ 1

#### General effect of small *ν*

The case of a small but finite *ν* is a singular pertubation of the *ν* = 0 system. We can investigate its behaviour using asymptotic methods in the small-*ν* limit. To briefly summarise before going into details, the main differences between the small- and the zero-*ν* cases is that small-*ν* dynamics do not vanish in region 𝒜, nor do they terminate at the point at which they first hit the region 𝒜-ℬ boundary. Instead, in the small-*ν* case the trajectories travel within region 𝒜 and along the region 𝒜-ℬ boundary over slow timescales of *O*(1*/ν*). So, in the small-*ν* case a trajectory starting in region 𝒞 will follow the zero-*ν* trajectory determined in Sec. 4.2 with an *O*(*ν*) correction, and move into region ℬ and then to the region 𝒜-ℬ boundary in finite time. A trajectory starting in region 𝒜 will also move into region ℬ over a slower time of *O*(1*/ν*). Given this, we focus our analysis on understanding what happens in region ℬ.

#### Asymptotic expansion

To understand the system behaviour in the limit of small *ν*, we first write both *y* and *α* as asymptotic series in powers of *ν*. That is *y*(*t*) ∼ *y*_0_(*t*) + *νy*_1_(*t*) and *α*(*t*) ∼ *α*_0_(*t*) + *να*_1_(*t*). Substituting these into the governing equations (10) and considering the leading-order equations, we obtain exactly the case *ν* = 0 that we analysed in Sec. 4.2. In this case, we know that 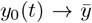 and 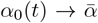 in some finite time *T*, where 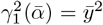. In the case where *ν* = 0, the analysis ends here. However, in the singular case of small but finite *ν*, there is an additional slow timescale after this finite time. To analyse this slow timescale behaviour, we transform into the slow timescale regime defined by *t* = *T* + *τ/ν*, where *τ* = *O*(1), and now expand *y*(*t*) = *Y* (*τ*) ∼ *Y*_0_(*τ*)+*νY*_1_(*τ*) and *α*(*t*) = *A*(*τ*) ∼ *A*_0_(*τ*)+*νA*_1_(*τ*). On substituting the slow timescale and these new expansions into the governing equations (10), we obtain

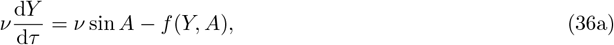

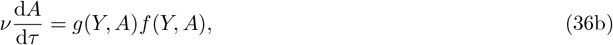

where we introduce the following shorthand functions

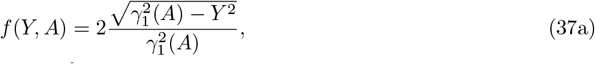

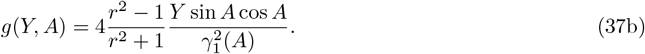

The *O*(1) terms in equations (36) generate a duplication of information, with both giving *f* (*Y*_0_, *A*_0_) = 0, or equivalently

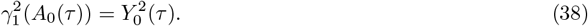

This tells us that the leading-order slow-time dynamics are confined to the line 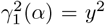, the boundary between regions 𝒜 and ℬ. The remaining goal of our analysis is to determine the precise dynamics of this slow motion. There are several ways to proceed at this point. One method involves continuing to the next asymptotic order and deriving an appropriate solvability condition. Another way involves combining the full equations (36) in a way that removes the duplication of information, then taking the limit *ν* → 0 to obtain an independent evolution equation. We proceed via the latter, since this involves significantly less algebra. By substituting (36a) into (36b) and dividing through by *ν*, we obtain

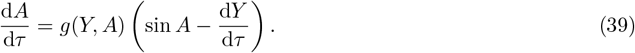

The equation (39) has leading-order form

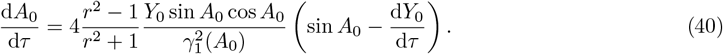

The equations (38) and (40) represent enough information to determine the slow evolution along the region 𝒜-ℬ boundary. It is more straightforward to see this if we use the direct time derivative of 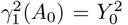 which is

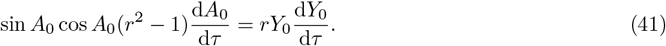

Combining this with (40) we can deduce that

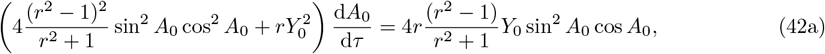

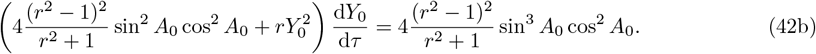

Then, since we are considering *α* ∈ [0, *π/*2] and therefore *A*_0_ ∈ [0, *π/*2], we define *s*_0_(*τ*) = sin(*A*_0_(*τ*)), where *s*_0_ ∈ [0, 1]. Under this substitution, (42) can be rewritten as

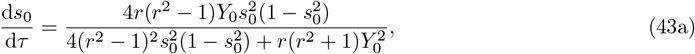

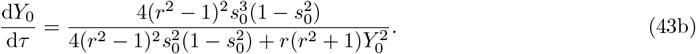

Solving (43) numerically, we can compare our numerical solution of the reduced problem to a numerical solution of the full problem. In Fig. 9, we see that the asymptotic solution gives good agreement with the full numerical solution for *ν* = 0.1, solved using `ode15s` in MATLAB. We also see that for all initial conditions, the trajectory will move to the point 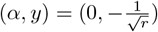, corresponding to perfect alignment.

**Figure 9:**
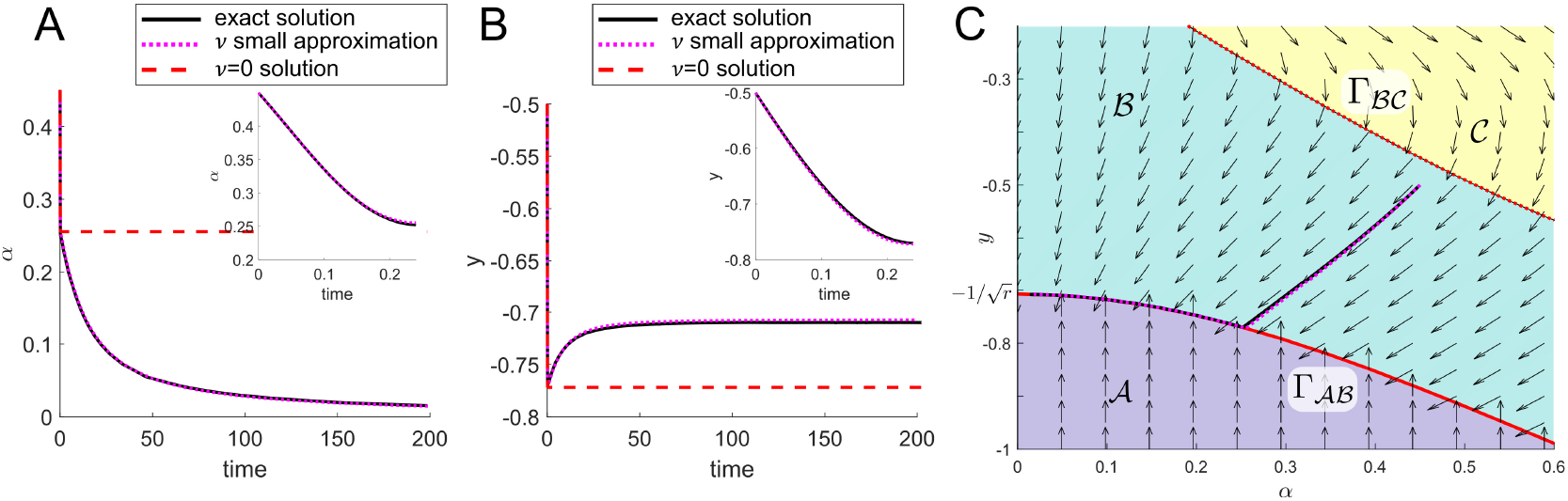
A, B: Comparison of *α*(*t*) (A) and *y*(*t*) (B) for solutions of the full system (solid-back), solutions for *ν* = 0 (dashed-red) and the small-*ν* approximation (43) (dotted-pink). Insets show dynamic for *t* small. C: Zoomed in region of phase portrait showing corresponding trajectory for A and B. Parameters *ν* = 0.1, *r* = 2.

If we substitute the solution of the algebraic constraint (38), 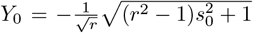 into (43a) we can reduce the dynamics to a single ODE

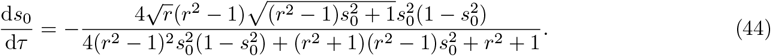

On separating variables and integrating, we find that

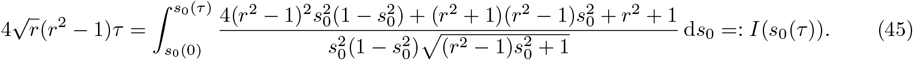

We note that we have shown in Sec. 4 that the point 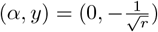 is a stable steady state for *α >* 0. We also know that all trajectories solving (43) move towards this point. To understand this long-term behaviour in more detail, we must understand what happens as *α* → 0^+^ or equivalently 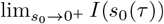. If this integral converges, then perfect alignment in reached in finite time. However, if this integral diverges, then it takes infinite time to reach perfect alignment where 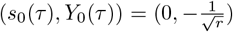 As *s*_0_ → 0, the integrand in (45) behaves like 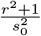 and hence the integral *I*(*s*_*0*_ (*τ*)) diverges. Therefore, the point 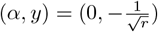 of perfect alignment is only reached in infinite time.

### 4.4 Lessons for (collective) cell alignment

#### The role of *ν*

The analysis of this section allows us to obtain a full understanding of the effect of *ν* on the alignment mechanism between two cells. As a reminder, *ν* was defined in (4) as 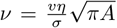 and can be interpreted as the ratio of self-propulsion and overlap avoidance strength in the presence of friction. We find that for *ν* = 0 trajectories do not move towards the point 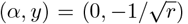, and hence perfect alignment is not achieved when there is no self-propulsion. This highlights the importance of self-propulsion in the alignment mechanism and shows that both overlap avoidance and self-propulsion are needed for full alignment. The analysis of 0 *< ν* ≪ 1 represents a singular perturbation of the *ν* = 0 case. We find that there is a set of initial conditions that lead to perfect alignment (for small, but positive *ν*), however, achieving perfect alignment takes infinite time. Quantifying computationally how favourable different model parameters are to alignment, we find a surprisingly good qualitative agreement with the results for large collective of cells found in Leech et al. (2024): *ν* = 0 leads to very little alignment, however, for larger values of *ν* alignment strength decreases as *ν* increases. The fact that the analysis for a symmetric two-cell situation recapitulates the results for large cell collectives indicates that it is a property of the alignment mechanism itself, not a result of the many-cell situation. Based on our analysis we can conclude that, while it is the overlap avoidance that causes interacting cells to align, (smaller amounts of) self-propulsion ensure that these interactions do not stop too quickly. However, if self-propulsion is too strong (compared to overlap avoidance), cells can “escape” the interaction by being pushed past each other.

#### The role of the aspect ratio *r*

For the dependence of alignment strength on the aspect ratio *r*, we found that for the symmetric two-cell system, there is an optimal aspect ratio for alignment: Interactions between close to circular cells will not lead to much velocity alignment. On the other hand, interactions between very elongated cells can more easily lead to cells moving past each other at which point the interaction stops. Note that this is different to the results for collectives found in Leech et al. (2024): There larger aspect ratios lead to more collective alignment. This could be, because in a collective, cell movement past each other might be hindered by other cells. Another possible reason could be, that *alignment* in Leech et al. (2024) referred to “nematic alignment”, which also captures situations where cells are side-by-side, but move in opposite directions. This is not something we can describe well with the symmetry condition imposed in this work, where *alignment* only describes velocity alignment, i.e. cells move in the same directions. A different framework would be necessary for further analytical investigations.

## 5 Discussion

### 5.1 Summary of the results

#### Model summary

In this work, we have presented a number of results to help us understand the alignment mechanism between two interacting cells in the model presented in Leech et al. (2024). These results have led to a more in-depth understanding of how model ingredients lead to alignment. Specifically we investigated the interplay between self-propulsion and overlap avoidance and the role of the aspect ratio of the cell. The analysed model describes self-propulsion and overlap avoidance, in reaction to which cells change their position, orientation and aspect ratio. We derived an analytical framework which led to a dynamical system, for which there are many mathematical tools available.

#### Deformable cells

In the full 3D (in variable space) system, which allows for shape deformations, a classic linear stability analysis is degenerate. Performing a dynamical asymptotic analysis, we find that the situation where both cells have their preferred shape and are perfectly aligned side-by-side, is a half-stable steady state of the system for elongated cells, and an unstable steady state for wide cells, indicating the fleeting nature of perfect alignment. This stability analysis also gives us insight into the timescales at play during the cell interactions. We find that the relative sizes of the two non-dimensional parameters determines the ordering of the change in aspect ratio and position, and that the change in orientation always occurs last, over a much slower timescale. We find that this change in orientation is unaffected by the shape change parameter, suggesting that deformability is not essential for alignment to occur. In the computational analysis done in Leech et al. (2024), it is found that allowing for changes in aspect ratio leads to greater alignment. The fact that we do not see this in the two-cell analysis, suggests that this result is due to the collective behaviour of the system, as opposed to the underlying alignment mechanism. We also find that if the non-dimensional shape change parameter is small, cells will quickly restore their preferred aspect ratio, after which the system behaves as though cell shape is rigid. This is reflected in the computational results in Leech et al. (2024) by noting that a strong shape-restoring force leads to little change in behaviour from the rigid cell case.

#### Non-deformable cells

We then consider a 2D (in variable space) limit of the full system in more detail, specifically, the limit in which the cells are rigid and keep their preferred aspect ratio. We computationally quantify the effect of model parameters on the overall alignment properties of the system. We find that both no self-propulsion *ν* = 0 and too much self-propulsion lead to less alignment, in striking qualitative agreement with the computation results in Leech et al. (2024) for cell collectives. This underlines the difference between active matter, that self-propels, and passive matter, which doesn’t. We are also able to make a significant amount of analytic progress in understanding the singular nature of the small self-propulsion limit. We find that cells are quickly pushed to a single point of overlap over a fast timescale, before the small self-propulsion causes cells to align while maintaining a single point of overlap over a much slower timescale. Together with the quantification of alignment, this leads to a greater understanding of how overlap avoidance combined with self-propulsion leads to alignment, see the discussion in Sec. 4.4. Hence, even though many of the complex details of the collective behaviour in Leech et al. (2024) are not captured in the analysis for two cells here, our analysis gives a plausible explanations for the observed behaviour. When analysing how alignment properties depend on the aspect ratio, we find that for our symmetric two-cell system there is an optimal aspect ratio for alignment. This is in contrast to the collective results in Leech et al. (2024), where larger aspect ratios lead to more alignment. This difference could be due to limitations of this work due to the imposed symmetry condition, which limit the notion of alignment to “velocity alignment”.

### 5.2 Further work

The modelling framework presented in Leech et al. (2024) is not specific to ellipse-shaped cells and could be applied to a number of different cell shapes. Provided the points of intersection could be found analytically, a similar approach to analyse and understand the system in greater depth could be taken. It would be interesting to apply the same analytical framework to different cell shapes to see how the results differ. This would be of particular interest if we wanted to apply the modelling framework to bacteria, for example, which are often modelled as spherocylinders instead of ellipses (Volfson et al., 2008; You et al., 2021). In this work we did not deeply analyse the behaviour of the system for wide cells with 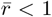, such as keratocytes. In this case a different symmetry condition might be appropriate to capture alignment: The two cells could be assumed to move one behind the other. In Leech et al. (2024) we also considered cell-cell adhesions. The analysis presented here could be extended to include such cell-cell contacts. In a first approximation, one could assume such contacts are permanent to then minimise a combined 2-cell energy. This will be subject of future work. Our approach could also be used to understand the dynamics of more than two cells, e.g. three cells placed in such a way that their centres of mass form a triangle. Finally, looking to other methods of mathematically analysing agent-based models, it would be beneficial to see whether any progress could be made on coarse-graining the model to obtain an equivalent continuum model such that further results on the collective behaviour of the system for many cells could be obtained. In the recent work of Merino-Aceituno et al. (2024) this is done for a similar model, which uses a smooth overlap potential.

## Acknowledgements

This work was supported by the Engineering and Physical Sciences Research Council (grant numbers EP/N509577/1 to VL and EP/T517793/1 to VL) which paid the salary of VL.

## Appendix

### A.1 Full stability analysis for *γ* = *O*(1)

Here we go through the full stability analysis of the steady state 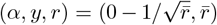 for *γ* = *O*(1). This is outlined briefly in the main text. We begin by perturbing the point 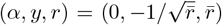 by a small amount, defined by

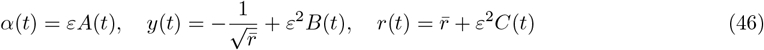

Understanding how *A*(*t*), *B*(*t*) and *C*(*t*) change over time will determine whether the point is stable or unstable. Under the conditions *ε* ≪ *γ* ≪ 1*/ε* with *ν* = *O*(1), that are three distinguished timescales of interest in the system: early time (*t* = *O*(*ε*)); intermediate time (*t* = *O*(*γ*)); and late time (*t* = *O*(1*/ε*)). Over the early timescale, *A*(*t*) remains unchanged, and *B*(*t*) and *C*(*t*) are driven by overlap avoidance. Over the intermediate timescale, *A*(*t*) still remains unchanged, *B*(*t*) and *C*(*t*) are still driven by overlap avoidance, and a restorative force restoring the aspect ratio to 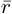 comes into the equation for *C*. Over the late timescale, the dynamics of *A*(*t*) become important and are driven by overlap avoidance. We see that the perturbations decay to 0 over the late timescale, proving that the steady state is stable. On substituting (46) into (10) we obtain

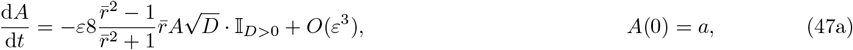

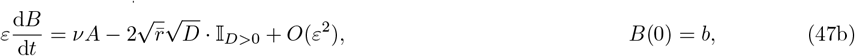

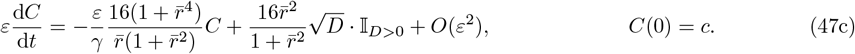

where 𝕀_*D>*0_ = 1 for *D >* 0 and zero otherwise, and *D* defined as

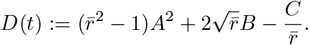

The sign of *D* determines whether we are in region 𝒜 or region ℬ and and hence whether overlap avoidance takes effect. We now analyse the system (47), starting with the early time.

#### A.1.1 Early time

We start our analysis over the early time, defined via *t* = *ετ* for *τ* = *O*(1). We indicate the early timescale variables with hats and write

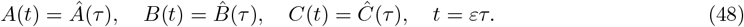

We obtain the following leading order equations by substituting (48) into (10).

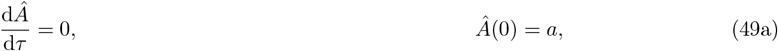

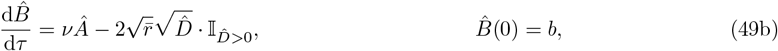

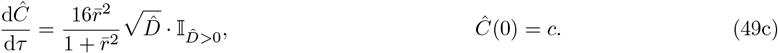

where 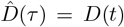. As in Sec. 3 (49a) shows that *Â*(*τ*) = *a* over the early time, and hence that the orientation remains constant over this timescale. Deriving an equation for 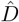 yields

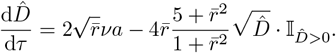

It is easy to see that if 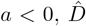 will become negative in finite time. Hence, even if cells interact initially, they will eventually stop interacting and move apart. In this case the steady state is not stable. If on the other hand 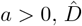 will become positive in finite time (even if it was negative initially) and convergence to a positive limiting value. In the following we will therefore focus on *a >* 0 and assume that cells interact. The first term in equation (49b) shows that self-propulsion will lead 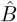 to increase over time. The second term in (49b) represents overlap avoidance, suggesting that cells will move apart to avoid overlap and consequently 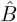 will decrease. The size of these two terms will determine whether 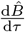 is positive or negative. 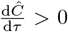 in (49c) means that the cells elongate over this timescale. This is because the orientation is near zero, so the cells are side-by-side and consequently elongate to avoid overlap.

##### Early time limiting behaviour

From (49) we can solve (49a) directly, and then combine (49b) and (49c) and integrate to obtain

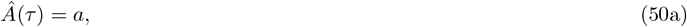

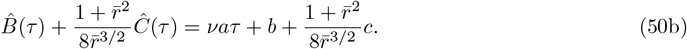

Equation (50b) shows that 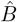 and 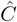 both grow linearly in time. We also find that, while 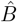 and 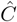 grow in time, the quantity

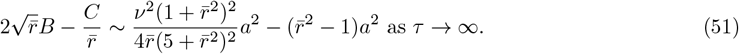

Specifically, this means that 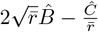 tends to a constant as *τ* grows. Combining (51), with the algebraic constraint for 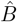 and 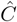 in (50b) we are able to solve for 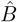 and 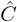 (as *τ* → ∞) and deduce that

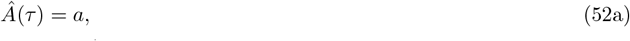

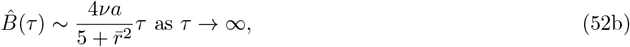

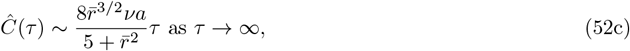

The fact that 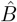 and 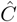 grow in time indicates the scaling for the next time region

#### A.1.2 Intermediate time

We now consider the intermediate timescale defined via 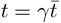, where 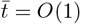, recalling that *ε* ≪ *γ* ≪ 1*/ε*. We also redefine our scalings for *B* and *C* as

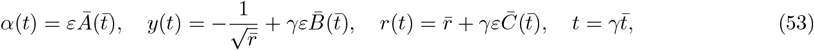

motivated by the large-*τ* results (52) in the early time in Sec. A.1.1. Substituting (53) into the governing equations (10) we obtain

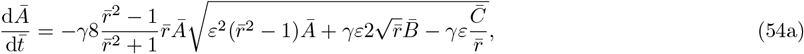

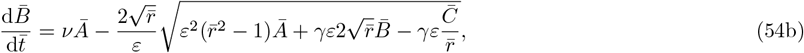

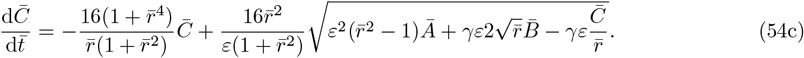

To leading order we obtain

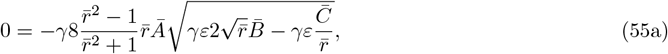

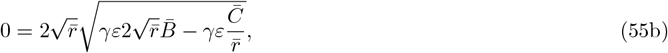

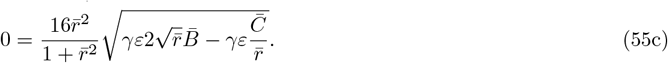

Solving (55) we see that we only obtain one piece of information:

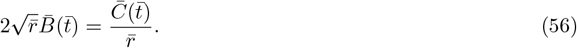

To obtain the additional information required, a formal analysis would involve expanding 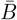 and 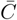 in higher powers of *ε*, and deriving appropriate solvability conditions. However, we can circumvent this laborious calculation by instead combining the equations in (54) to remove the duplication of information at leading order (essentially the square root term in each equation), and generate a system of distinct ODEs for 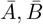 and 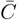 We can eliminate the square root term in (54a), (54b) and (54c) to obtain

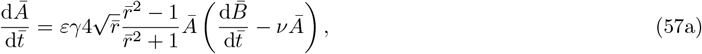

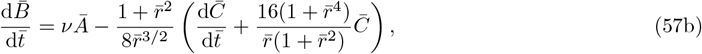

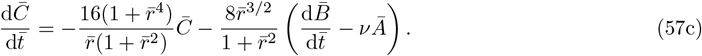

We can then decouple (57b) and (57c) using (56). On doing this, to leading order we obtain

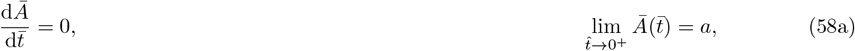

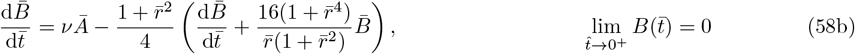

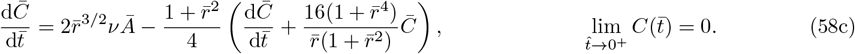

The initial conditions are matching conditions from the previously solved early timescale *τ*. Solving (58) we obtain

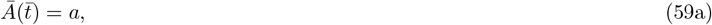

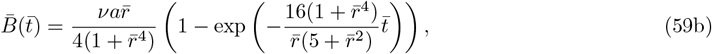

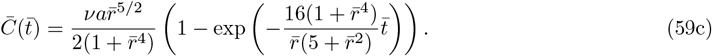

From (59) we see that, again, *Ā*remains constant over the intermediate time, and that 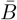 and 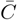 decay exponentially in time, reaching constant values given by

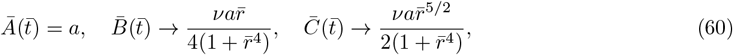

as 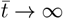 These will be used as matching conditions for the next timescale.

#### A.1.3 Late time

Finally, we consider the late timescale defined via 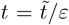, where 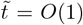 Given the large-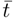 results (60), we retain the scalings from the intermediate time region in (53).

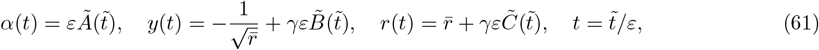

This is the timescale over which the dynamics of *A* become important. Over this timescale equations (10) become

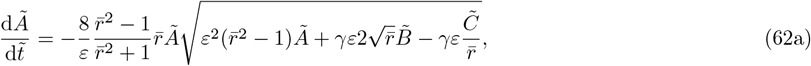

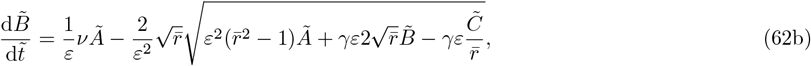

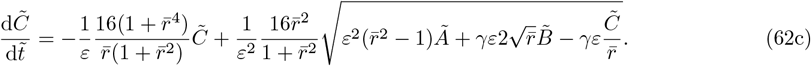

To leading order we obtain

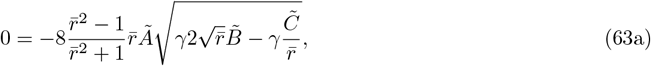

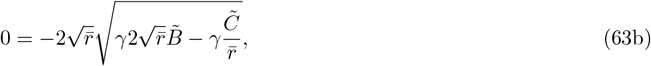

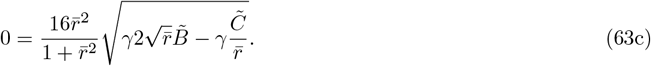

Again, we obtain only one piece of information:

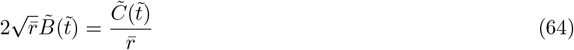

Since this information is duplicated in all three equations, we would have to expand 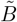and 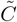 in higher orders of *ε* to obtain solvability conditions. Alternatively, as in the intermediate time region, we can rearrange the equations to eliminate the square root terms and generate a system of ODEs for 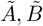 and 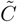. On doing so we obtain

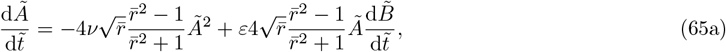

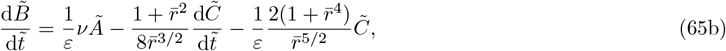

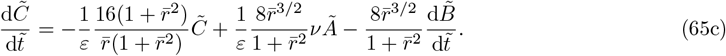

To leading order we have

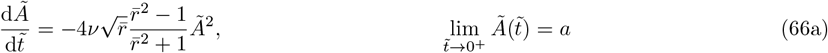

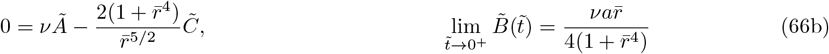

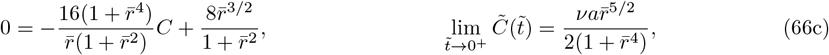

where the matching conditions come from the long time behaviour (60) in the previous timescale. Solving (66) using (64) we find that

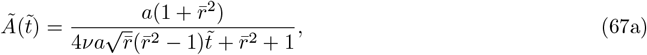

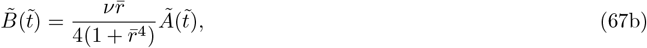

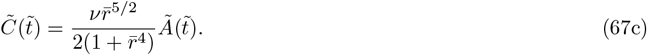

For 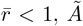, *Ã* will blow up in finite time and hence the steady state is unstable. For 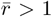 (and *a >* 0) on the other hand, we can see that 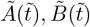 and 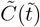 all decay to zero algebraically as 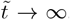 This shows that the point we are interested in is half-stable for 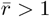. We note that this analysis for *γ* = *O*(1) is just a splitting of the early timescale in the analysis done in the main text.

## References

Albi, G. and Pareschi, L. 2013. Modeling of self-organized systems interacting with a few individuals: from microscopic to macroscopic dynamics. Appl. Math. Lett., 26(4):397–401.

Balagam, R. and Igoshin, O. A. 2015. Mechanism for collective cell alignment in myxococcus xanthus bacteria. PLoS Comput. Biol., 11(8):e1004474.

Baskaran, A. and Marchetti, M. C. 2008. Enhanced diffusion and ordering of self-propelled rods. Phys. Rev. Lett., 101(26):268101.

Bolley, F., Cañizo, J. A., and Carrillo, J. A. 2012. Mean-field limit for the stochastic vicsek model. Appl. Math. Lett., 25(3):339–343.

Dartsch, P. and Betz, E. 1989. Response of cultured endothelial cells to mechanical stimulation. Basic Res. Cardiol., 84(3):268–281.

Degond, P. and Motsch, S. 2008. Continuum limit of self-driven particles with orientation interaction. Math. Models Methods Appl. Sci., 18(supp01):1193–1215.

Deutsch, A. and Dormann, S. 2005. Mathematical modeling of biological pattern formation. Springer, New York.

Erdogan, B., Ao, M., White, L. M., Means, A. L., Brewer, B. M., Yang, L., Washington, M. K., Shi, C., Franco, O. E., Weaver, A. M., et al. 2017. Cancer-associated fibroblasts promote directional cancer cell migration by aligning fibronectin. J. Cell Bio., 216(11):3799–3816.

Fay, T. H. and Joubert, S. V. 2010. Separatrices. Int. J. Math. Educ. Sci. Technol., 41(3):412–418.

Großmann, R., Peruani, F., and Bär, M. 2016. Mesoscale pattern formation of self-propelled rods with velocity reversal. Phys. Rev. E, 94(5):050602.

Herbert-Read, J. E., Perna, A., Mann, R. P., Schaerf, T. M., Sumpter, D. J., and Ward, A. J. 2011. Inferring the rules of interaction of shoaling fish. Proc. Natl. Acad. Sci., 108(46):18726–18731.

Kenny, F. N., Marcotti, S., De Freitas, D. B., Drudi, E. M., Leech, V., Bell, R. E., Easton, J., Fleck, R., Allison, L., Philippeos, C., et al. 2023. Autocrine il-6 drives cell and extracellular matrix anisotropy in scar fibroblasts. Matrix Biol., 123:1–16.

Kraikivski, P., Lipowsky, R., and Kierfeld, J. 2006. Enhanced ordering of interacting filaments by molecular motors. Phys. Rev. Lett., 96(25):258103.

Leech, V., Kenny, F. N., Marcotti, S., Shaw, T. J., Stramer, B. M., and Manhart, A. 2024. Derivation and simulation of a computational model of active cell populations: How overlap avoidance, deformability, cell-cell junctions and cytoskeletal forces affect alignment. PLOS Computational Biology, 20(7):e1011879.

Lopez, U., Gautrais, J., Couzin, I. D., and Theraulaz, G. 2012. From behavioural analyses to models of collective motion in fish schools. Interface Focus, 2(6):693–707.

Merino-Aceituno, S., Plunder, S., Wytrzens, C., and Yoldaş, H. 2024. Macroscopic effects of an anisotropic gaussian-type repulsive potential: nematic alignment and spatial effects. arXiv preprint 2410.06740.

Ng, C. P. and Swartz, M. A. 2003. Fibroblast alignment under interstitial fluid flow using a novel 3-d tissue culture model. American Journal of Physiology-Heart and Circulatory Physiology, 284(5):H1771–H1777.

Peruani, F., Deutsch, A., and Bär, M. 2006. Nonequilibrium clustering of self-propelled rods. Phys. Rev. E. Stat. Nonlin. Soft Matter Phys., 74(3):030904.

Van Dyke, M. 1975. Perturbation methods in fluid mechanics. Stanford, CA: Parabolic Press, California.

Volfson, D., Cookson, S., Hasty, J., and Tsimring, L. S. 2008. Biomechanical ordering of dense cell populations. Proc. Natl. Acad. Sci., 105(40):15346–15351.

Wang, W. Y., Pearson, A. T., Kutys, M. L., Choi, C. K., Wozniak, M. A., Baker, B. M., and Chen, C. S. 2018. Extracellular matrix alignment dictates the organization of focal adhesions and directs uniaxial cell migration. APL Bioeng, 2(4).

You, Z., Pearce, D. J., and Giomi, L. 2021. Confinement-induced self-organization in growing bacterial colonies. Sci. Adv., 7(4):eabc8685.

